# Dendritic cells require TMEM176A/B ion channels for optimal MHC II antigen presentation to naive CD4^+^ T cells

**DOI:** 10.1101/851527

**Authors:** Melanie Lancien, Geraldine Bienvenu, Lucile Gueno, Sonia Salle, Emmanuel Merieau, Severine Remy, Amandine Even, Aurelie Moreau, Alice Molle, Cynthia Fourgeux, Flora Coulon, Gaelle Beriou, Laurence Bouchet-Delbos, Elise Chiffoleau, Peggy Kirstetter, Susan Chan, Steven Kerfoot, Saeed Abdu Rahiman, Veronica De Simone, Gianluca Matteoli, Gaelle Boncompain, Franck Perez, Regis Josien, Jeremie Poschmann, Maria Cristina Cuturi, Cedric Louvet

**Affiliations:** Nantes Université, CHU Nantes, Inserm, Centre de Recherche en Transplantation et Immunologie, UMR 1064, ITUN, F-44000 Nantes, France; Institut de Génétique et de Biologie Moléculaire et Cellulaire (IGBMC), INSERM U1258, CNRS UMR 7104, Université de Strasbourg, Illkirch, France; Department of Microbiology and Immunology, University of Western Ontario, London, Ontario N6A 5C1, Canada; Department of Chronic Diseases, Metabolism and Ageing, Translational Research Center for Gastrointestinal Disorders (TARGID), University of Leuven, Leuven, Belgium; Dynamics of Intracellular Organization Laboratory, Institut Curie, PSL Research University, Sorbonne Université, Centre National de la Recherche Scientifique, UMR 144, Paris, France

## Abstract

Intracellular ion fluxes emerge as critical actors of immunoregulation but still remain poorly explored. Here we investigated the role of the redundant cation channels TMEM176A and TMEM176B (TMEM176A/B) in RORγt^+^ cells and conventional dendritic cells (cDCs) using germline and conditional double knock-out (DKO) mice. While *Tmem176a/b* appeared surprisingly dispensable for the protective function of Th17 and group 3 innate lymphoid cells (ILC3s) in the intestinal mucosa, we found that they were required in cDCs for optimal antigen processing and presentation to CD4^+^ T cells. Using a real-time imaging method, we show that TMEM176A/B accumulate in dynamic post-Golgi vesicles preferentially linked to the late endolysosomal system and strongly colocalize with HLA-DM. Together, our results suggest that TMEM176A/B ion channels play a direct role in the MHC II compartment (MIIC) of DCs for the fine regulation of antigen presentation and naive CD4^+^ T cell priming.

## Introduction

Multiple ion channels and transporters are expressed in both innate and adaptive immune cells to control various vital functions, from membrane potential regulation to receptor signaling or migration^1^. The role of ion flux has notably been fully appreciated following the molecular characterization of the store-operated Ca^2+^ entry (SOCE) through Ca^2+^ release–activated Ca^2+^ (CRAC) channels mediated by the ORAI/STIM complex, best characterized in T cells. However, this system appears dispensable for key functions of macrophages and dendritic cells (DCs) while Ca^2+^ signaling remains critical in these cells^2,3^. This observation points to the importance of alternative systems, yet to be discovered, that regulate the intracellular and luminal concentrations of Ca^2+^ but also other ions including Na^+^, K^+^, Cl^−^, Mg^2+^ or Zn^2+^ for the control of immune responses. In comparison with the plasma membrane, there is still a paucity of studies investigating the role of intracellular ion channels and transporters, notably in DCs^4^, that could provide major insights into the understanding of new immunomodulatory mechanisms.

We previously showed that the co-regulated genes *Tmem176a* and *Tmem176b* encode redundant acid-sensitive, non-selective, cation channels^5,6^ whose precise functions remain largely unknown in vivo. Transcriptomic, SNP or epigenetic analysis have associated these homolog genes with different pathologies such as multiple sclerosis^7^, chronic obstructive pulmonary disease (COPD)^8^ or age-related macular degeneration (AMD)^9^. These findings suggest an important role of *Tmem176a/b* in the development of inflammatory diseases thus emphasizing the need to identify the immune cell types and the functions in which they are predominantly involved.

We initially cloned *Tmem176b* (originally named *Torid*) as an over-expressed gene encoding an intracellular four-span transmembrane protein in myeloid cells infiltrating non-rejecting allografts^10^. We later demonstrated its contribution to the suppressive function of ex vivo-generated tolerogenic DCs through antigen cross-presentation by allowing cation (Na^+^) counterflux required for progressive endophagosomal acidification^5^. However, the function of this ion flux in the homeostasis and physiological response of conventional DCs (cDCs) has not been explored and is likely achieved by both TMEM176A and TMEM176B in a redundant fashion^6^.

Unexpectedly, besides myeloid cells, we and others reported a strong expression of *Tmem176a* and *Tmem176b* in the retinoic-acid-receptor-related orphan receptor-γt-positive (RORγt^+^) lymphoid cell family, also referred to as type 3 (or type 17) immune cells, producing the prototypical cytokines IL-17 and IL-22 and including Th17 CD4^+^ T cells, γδT17 cells, group 3 innate lymphoid cells (ILC3s) and NKT17 cells^6,11–14^. Moreover, the Littman group included *Tmem176a/b* in the restricted group of 11 genes whose expression is directly dependent on RORγt in Th17 cells^15^. However, in our previous study, we found no or only modest effect of *Tmem176b* single deficiency in different models of autoinflammation linked to type 3 immunity^6^. We speculated that the absence of *Tmem176b* could be efficiently compensated by its homolog *Tmem176a*, located in the same genomic locus, thus masking possible phenotypic alterations.

Here we have used germline and conditional (“floxed”) double knock-out (DKO) mice to unequivocally determine the importance of *Tmem176a* and *Tmem176b* in the biology of RORγt^+^ cells and cDCs in vivo. In that respect, this is the first study exploring the consequence of *Tmem176a/b* double deficiency in vivo. Furthermore, we have combined these functional results with the elucidation of the precise trafficking of both proteins using a real-time imaging method. Our findings show that, while *Tmem176a/b* appear surprisingly dispensable for RORγt^+^ cell functions, these genes are required in the MHC II pathway in DCs for efficient priming of naive CD4^+^ T cells.

## Results

### Generation of germline and conditional double KO (DKO) mice simultaneously targeting Tmem176a and Tmem176b

*Tmem176a* and *Tmem176b* are homolog genes encoding structurally similar four-span transmembrane proteins^6,10,16^ (**Figure 1A**). In the immune system, our previous studies^6,10,17^ combined with the analysis of Immuno-Navigator^18^ and Immgen^19^ public databases (**Figure S1**) indicate that *Tmem176a* and *Tmem176b* are tightly co-regulated and highly expressed both in cDCs and in the RORγt^+^ cell family (depicted in **Figure 1B**). Taken together, these observations along with reported evidence of genetic compensation in *Tmem176b*^−/−^ single KO mice^6^ strongly suggested the need to simultaneously target both genes to decipher their function.

**Figure 1.**
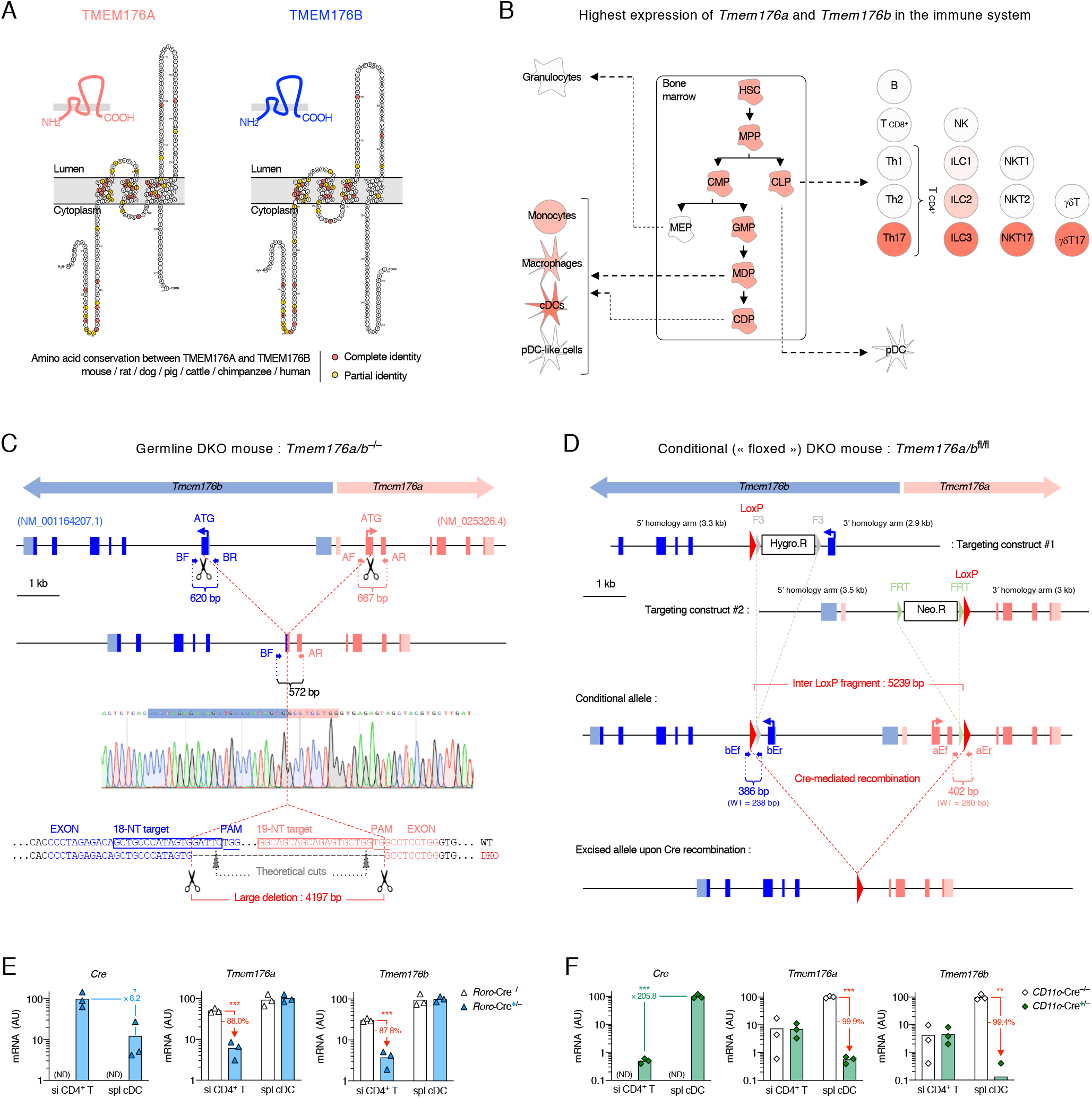
Generation of *Tmem176a/b* double-deficient mice. (**A**) Graphical topology of mouse TMEM176A and TMEM176B using Protter. The most conserved amino acids between both molecules and across multiple species are highlighted and mainly concentrate within the three first transmembrane domains and in the N-terminal region. (**B**) Synthetic view of *Tmem176a* and *Tmem176b* expression in immune cells summarized from the literature and public databases (Figure S1). Dark red represents the highest expression. (**C**) Generation of germline *Tmem176a/b* double KO (DKO) mice carrying a large deletion using a dual CRISPR–Cas9 strategy. *Tmem176a* and *Tmem176b* genes are oriented in an opposite direction in the same genomic locus and share analogous intron-exon organization, with the respective first coding exons only separated by 3.8 kb. Small arrows labeled BR, BR, AF and AR show the position of the primers used for PCR genotyping of the mutant mice. Exons are shown as filled boxes and untranslated regions as shaded boxes. (**D**) Generation of conditional *Tmem176a/b* DKO mice. Small arrows labeled bEf, bEr, aEf and aEr show the position of the primers used for PCR genotyping of the mutant mice. (**E**-**F**) Verification of cell-specific deletion in *Tmem176a/b* DKO conditional mice crossed to *Rorc*-Cre (**E**) or *CD11c*-Cre (**F**) mice. Relative mRNA expression of the indicated genes quantified by qPCR in purified small intestine (si) CD4^+^ T cells from mice treated with anti-CD3 and in purified splenic cDCs from naive mice.

As described previously in a methodological study^20^, we generated a germline double KO (DKO) mouse (**Figure 1C**) in the pure C57BL/6N genetic background. Homozygous DKO mice were born in a Mendelian ratio and appeared normal, without any significant difference in weight from control WT littermates, suggesting that *Tmem176a/b* are not absolutely required, or can be sufficiently compensated, during development.

In order to clearly associate a cell type with a potential phenotypic observation resulting from *Tmem176a/b* deficiency, we also engaged in the generation of a conditional DKO mouse (*Tmem176a/b*^flox^) (**Figure 1D**) that we crossed to *Rorc*(γt)-Cre and *CD11c*-Cre (*Itgax*-Cre) mice to target RORγt^+^ and CD11c^+^ cells, respectively. Verification of differential Cre-driven cell-specific deletion was performed by comparing CD4^+^ T cells from the small intestine (si) and splenic cDCs for their expression of Cre, *Tmem176a* and *Tmem176b* mRNA (**Figure 1E-F**). To increase the fraction of RORγt^+^ Th17 cells in siCD4^+^ T cells, mice were injected with anti-CD3 as previously described^21^. As anticipated, Cre expression was highest in siCD4^+^ T cells from *Rorc*-Cre mice and in cDCs from *CD11c*-Cre mice.

Importantly, both *Tmem176a* and *Tmem176b* transcriptional levels were specifically decreased in those subsets as compared to Cre-negative littermate mice. Although deletion efficiency appeared lower in siCD4^+^ T cells than in cDCs, this could be explained by the yet relatively low fraction of RORγt^+^ cells (61.9%±7.9) within our preparations of purified siCD4^+^ T cells. Finally, it is interesting to note that cDCs from *Rorc*-Cre mice exhibited a substantial level of Cre expression, a finding in line with a recent study^22^ describing a subset of cDC expressing *Rorc*, which however did not appear sufficient to allow Cre-mediated allele recombination.

### Tmem176a/b DKO mice exhibit normal ILC3 and Th17 distribution and RORγt cell-dependent protective functions in the gut mucosa

Owing to the high expression of *Tmem176a/b* in RORγt^+^ cells, we focused our attention on the intestinal mucosa where these cells preferentially localize to exert their sentinel function in response to the gut microbiota^23^. Flow cytometry analysis (gating strategy depicted in **Figure S2A-B**) of CD4^+^ T cells and ILCs in the small intestine and colon lamina propria revealed no differences in the proportions of Th17 or ILC3 subsets between WT and DKO mice (**Figure 2A**). Furthermore, in vitro cytokine production of intestinal CD4^+^ T cells and ILCs was not affected by *Tmem176a/b* double deficiency (**Figure 2B**). Concordant with this, we did not detect any significant change in the level of expression of *Rorc2* (the isoform encoding RORγt), *Il23r*, *Il22*, *Il17a* or *Il17f* as well as target genes of the IL-22/IL17 axis in the small intestine and colon of DKO mice (**Figure S2C**).

**Figure 2.**
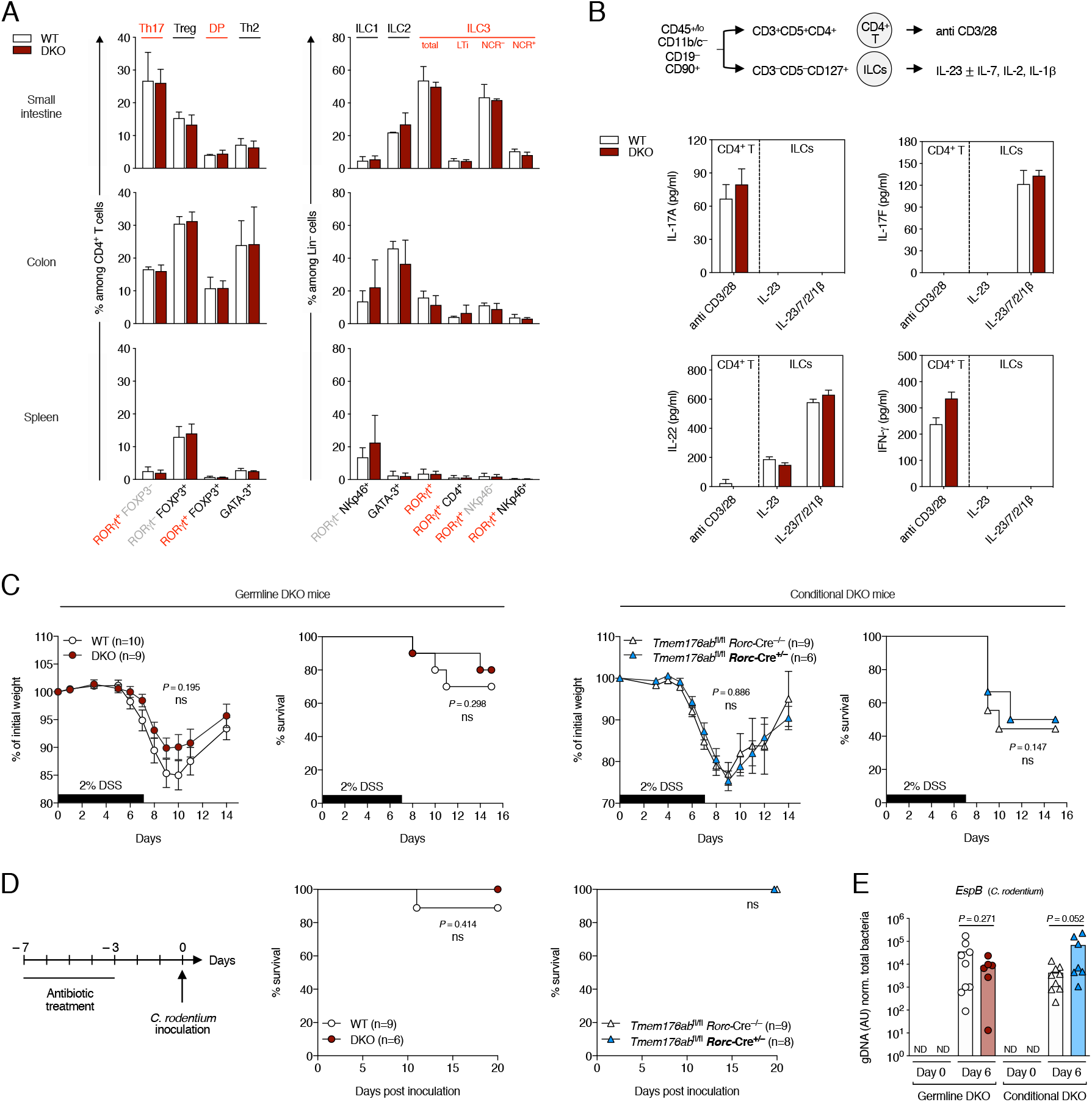
Th17 and ILC3 frequencies and RORγt^+^-cell protective functions in the gut mucosa of *Tmem176a/b* double-deficient mice. (**A**) Relative frequencies of the indicated populations in the small intestine (siLP), colon (cLP) lamina propria and spleen of WT and *Tmem176a/b* DKO littermate mice. Data shown are means (±SD) of three independent experiments. (**B**) In vitro cytokine secretion by CD4^+^ T cells and ILCs FACS-sorted from the intestinal lamina propria of WT and *Tmem176a/b* DKO littermate mice following in vitro culture (18 h) with the indicated stimuli. Data shown are means (±SD) of triplicates representative of three independent experiments. (**C**) Acute colitis induced with 2% DSS in drinking water for 7 consecutive days in germline *Tmem176a/b* (left panels) and RORγt^+^ cell-restricted conditional (right panels) DKO mice in comparison to control mice. Data are presented as percent of initial weight (±SEM) and survival (weight loss >20%). (**D**) Infection with *C. rodentium* administered orally (2×10^9^ CFU) in germline *Tmem176a/b* and RORγt^+^ cell-restricted conditional DKO mice in comparison to control mice. (**E**) Quantification of *C. rodentium EspB* gene by qPCR in the faeces prior infection and 6 days post-infection. Bars indicate means and dots represent individual mice. ND : Not detected.

To test whether *Tmem176a/b* could play a role in RORγt^+^ cells in the context of inflammatory responses in the gut mucosa, we used the injury-induced self-resolving model of dextran sodium sulfate (DSS)-induced acute colitis in which Th17/ILC3-derived IL-22 is pivotal to restore barrier integrity upon epithelial damage^24,25^. However, both germline DKO and *Tmem176a/b*^fl/fl^*Rorc*-Cre^+/–^ conditional mice exhibited a similar response to DSS when compared to control mice (**Figure 2C**). Next, we used an infectious colitis model using *Citrobacter rodentium*, a mouse attaching and effacing bacterial pathogen considered as an excellent model of clinically important human gastrointestinal pathogens and against which ILC3s and Th17 have been shown to play a protective and redundant role^11,26,27^. As shown in **Figure 2D**, *Tmem176a/b* deficiency did not hamper the mouse resistance to this infection. Moreover, although fecal *Citrobacter rodentium* bacterial loads detected by qPCR appeared increased in RORγt-specific conditional DKO, this difference did not reach statistical significance and was not found in germline DKO mice compared to respective control mice (**Figure 2E**).

Thus, these results suggest that *Tmem176a/b* are not critical for the development and protective functions of ILC3s and Th17 in the intestinal mucosa.

### DCs develop normally in Tmem176a/b-deficient mice

Next, we investigated the effect of *Tmem176a/b* double deficiency in the biology of cDCs. First, because both genes are highly expressed in the majority of hematopoietic precursors including cDC progenitors (**Figure 1B** and **Figure S1**), we assessed their frequency in the bone marrow but found no alteration in DKO mice (**Figure 3A** **and Figure S3**). Accordingly, the percentages and absolute numbers of CD11c^high^ MHC II^+^ cells in the spleen and, within this population, the proportions of the two main conventional DC subsets, namely cDC1 (CD11b^−^ CD8α^+^) and cDC2 (CD11b^+^ CD8α^−^), were similar between WT and DKO mice (**Figure 3B**). Furthermore, MHC I, MHC II, CD80 (B7-1) and CD86 (B7-2) surface expression on cDC1 and cDC2 were not affected by *Tmem176a/b* deficiency and LPS stimulation elicited an equally strong upregulation of MHC II, CD80 and CD86 at the surface of both WT and DKO cDC subsets (**Figure 3C**). Finally, purified spleen cDCs from WT and DKO mice produced similar basal levels of IL-12 and IL-6 that were both increased with the addition of LPS (**Figure 3D**).

**Figure 3.**
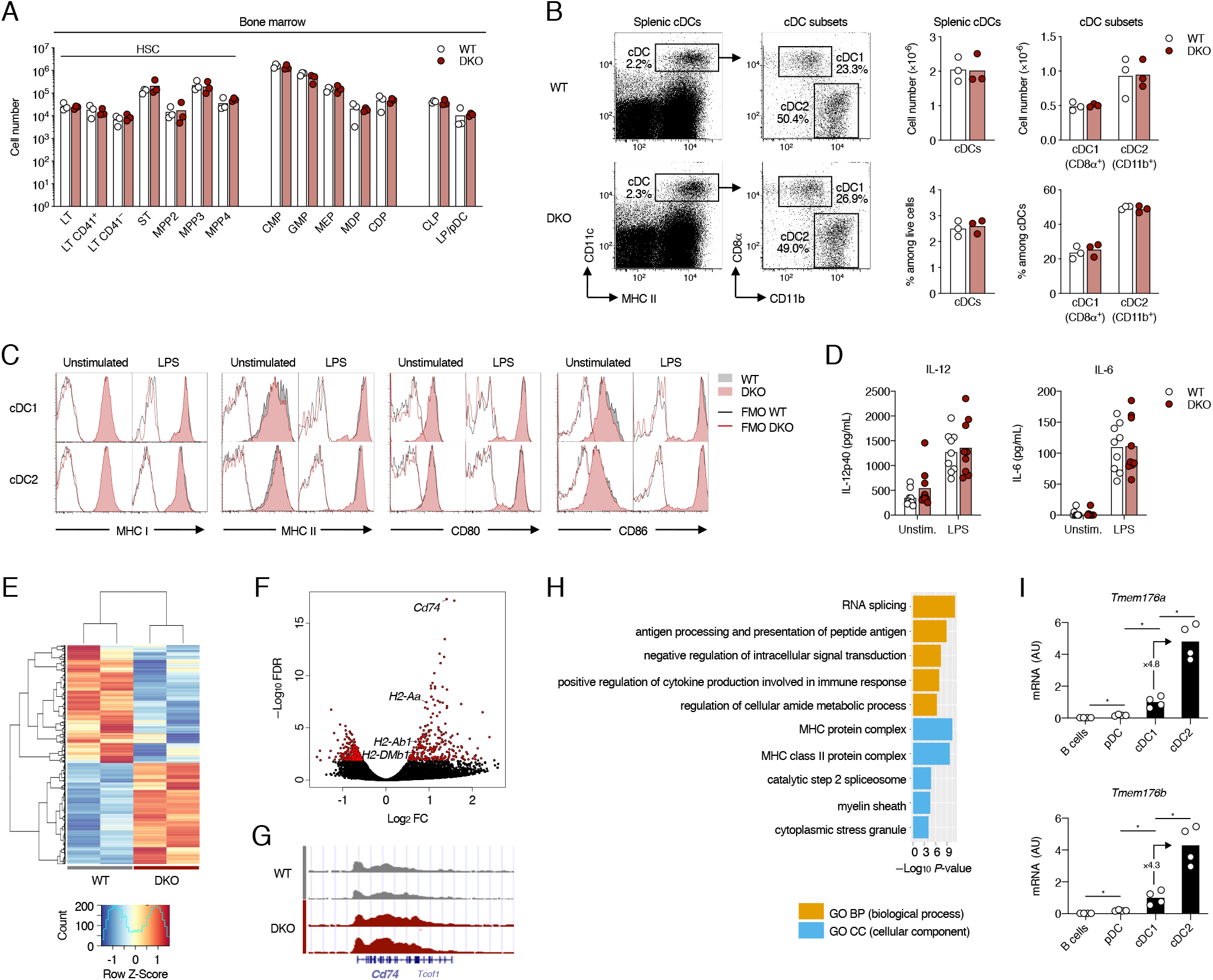
Phenotypic analysis of cDCs in *Tmem176a/b* DKO mice. (**A**) Hematopoietic stem cell and progenitor numbers in the bone marrow of WT and *Tmem176a/b* DKO mice analyzed by flow cytometry. Bars indicate means and dots represent individual mice. (**B**) Gating strategy (left panels) and quantification (right panels) of spleen cDCs and subsets from WT and *Tmem176a/b* DKO littermate mice by flow cytometry. Bars indicate means and dots represent individual mice. Data were pooled from three independent experiments. (**C**) Expression of MHC I, MHC II, CD80 and CD86 molecules at the surface of spleen cDC1 and cDC2 from WT and *Tmem176a/b* DKO littermate mice. Cells were analyzed by flow cytometry freshly after cell preparation or upon LPS stimulation for 6 h in vitro. Shown histograms data are representative of three independent experiments. FMO : Fluorescence minus one. (**D**) IL-12 and IL-6 secretion by purified cDCs purified from the spleen of WT and *Tmem176a/b* DKO littermate mice and cultured for 16 h with or without LPS. Data were pooled from three independent experiments. (**E**) Heatmap showing unsupervised clustering z-scores of the differentially acetylated (DA) peaks between WT and DKO spleen cDCs. (**F**) Volcano plot of the 962 DA peaks (red). Relevant genes linked to MHC class II pathway are highlighted. (**G**) Genome browser of *Cd74*, hyperacetylated in DKO. (**H**) Gene ontology of the 445 DA peaks up-regulated in DKO vs WT. (**I**) Expression of *Tmem176a* and *Tmem176b* assessed by RT-qPCR in the indicated populations FACS-sorted from the spleen. Bars indicate means and dots represent individual mice.

In the absence of obvious developmental abnormalities of cDCs, we sought to identify dysregulated cellular pathways resulting from the absence of *Tmem176a/b*. Chromatin dynamics reflect with great sensitivity gene regulation and play important roles in immune functions such as trained innate immunity^28^ or T cell effector/memory differentiation^29^. To investigate whether epigenetic alterations are present in DKO cDCs we FACS-sorted CD11c^high^MHC II^+^ cells and performed chromatin immunoprecipitation by targeting lysine H3K27 acetylation (H3K27ac) followed by sequencing (ChIP-seq) to detect active promoters and enhancers (epigenomics road map). We found that 962 enhancers were significantly differentially acetylated (DA) between WT and DKO cDCs (**Figure 3E-F**). Interestingly, one of the most significantly DA gene regulatory region covered the entire *Cd74* gene (**Figure 3G**) and MHC gene ontology analysis included antigen processing and MHC II protein molecules (**Figure 3H**). Together, these data suggest that *Tmem176a/b* deficiency could impact intracellular processes of the MHC II pathway and lead to selective genetic adaptations. Further supporting a role of *Tmem176a/b* in the MHC II pathway, we found that both homologs were clearly over-expressed in cDC2 when compared to B cells, pDCs and cDC1 (**Figure 3I**), consistent with the propensity of this subset in priming naive CD4^+^ T cells through MHC II antigen presentation^30^.

### Tmem176a/b are required for optimal presentation of exogenous antigens to CD4^+^ T cells by DCs

To determine whether *Tmem176a/b* are required in DCs for antigen presentation in vivo via MHC I or MHC II molecules, we set out to test different models of dominant CD8^+^ or CD4^+^ T cell responses. First, using a minor histocompatibility transplantation model in which male skin is grafted onto female recipients, we found that WT and DKO exhibited the same rate of graft rejection (**Figure 4A**), suggesting that CD8^+^ T cell priming and effector functions were not impaired in the absence of *Tmem176a/b*. Next, we examined whether anti-tumor immune responses could be influenced by *Tmem176a/b* deficiency. Notably, we hypothesized that DKO could exhibit enhanced anti-tumor immunity owing to a recent study proposing that targeting *Tmem176b* could improve anti-tumor CD8^+^ T-cell response by de-repressing inflammasome activation in myeloid cells^31^. Subcutaneous injection of OVA-expressing EG7 thymoma led to detectable tumors within two weeks in all mice and, consistently with the relatively immunogenic nature of this cell line in our experimental conditions, tumor regression was then observed in a large fraction of the mice but without significant difference between WT and DKO mice (**Figure 4B**). Furthermore, *Tmem176a/b* deficiency did not impact tumor growth and mouse survival with two aggressive tumor cell lines, MCA101-sOVA fibrosarcoma and B16-OVA melanoma (**Figure 4B**). Thus, our results show that the absence of *Tmem176a* and *Tmem176b* does not prevent nor enhance CD8^+^ T-cell responses. To evaluate CD4^+^ T-cell responses in vivo, we first used the model of experimental autoimmune-encephalomyelitis (EAE) induced by MOG_35–55_ peptide or MOG_1–125_ protein immunization (**Figure 4C**). Interestingly, whereas EAE developed similarly between WT and DKO mice with MOG peptide immunization (**Figure 4D**), DKO mice appeared less susceptible if the MOG protein was used (**Figure 4E**), thus pointing to a specific role of *Tmem176a/b* in antigen processing before peptide-MHC II complex display at the surface. Because of the high variability of this model, we next turned to a model of delayed-type hypersensitivity (DTH) (**Figure 4F**). Again, we observed that DKO mice differed from WT mice for the DTH response upon challenge in the footpad only if the whole protein was used for the immunization (**Figure 4G-H**). Taken together, these data suggest that *Tmem176a/b* deficiency selectively affects the intracellular processing of exogenous antigens for naive CD4^+^ T-cell priming.

**Figure 4.**
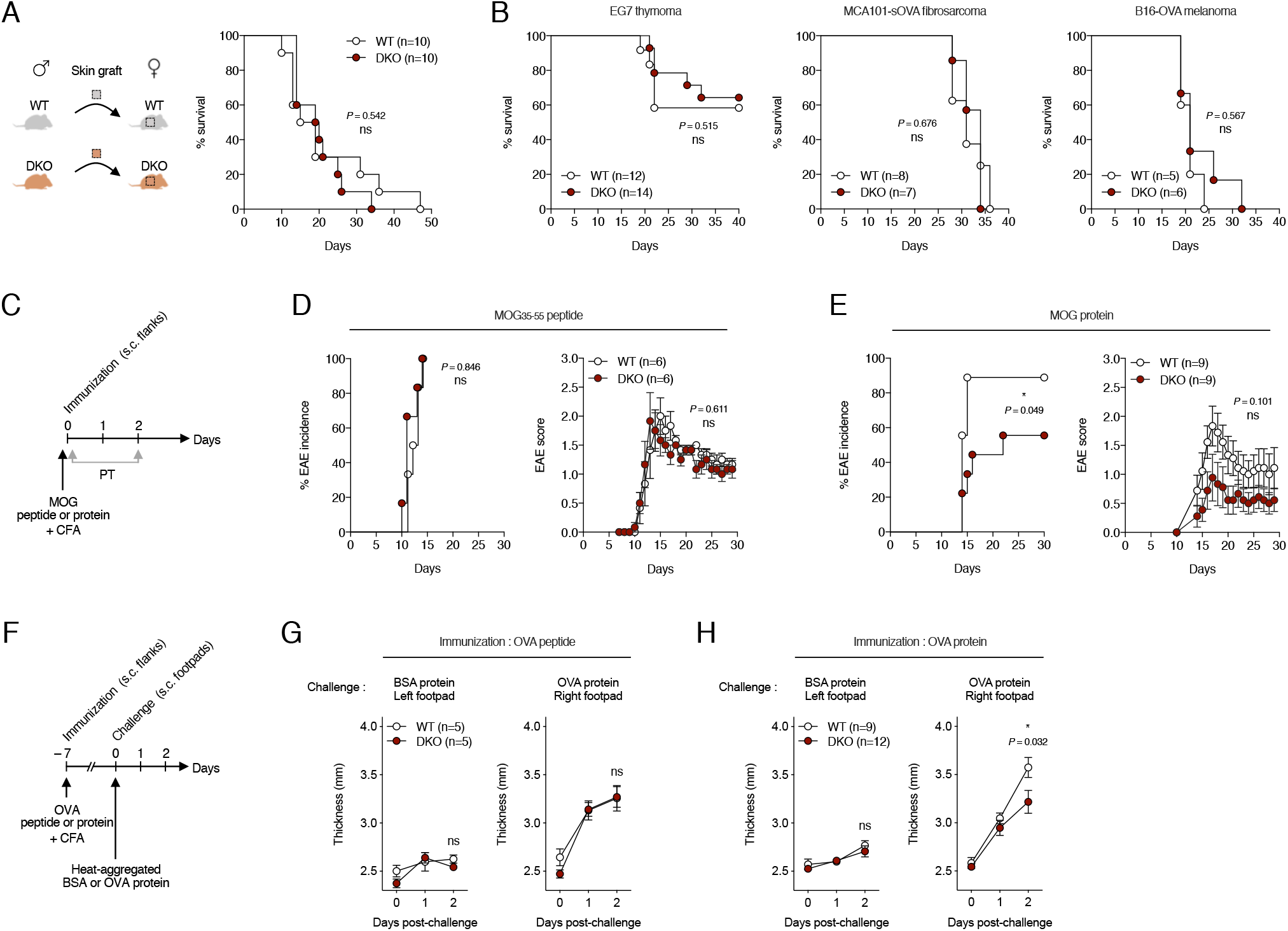
Evaluation of CD8^+^ and CD4^+^ T cell-dependent models in *Tmem176a/b* DKO mice. (**A**). Rejection rates of male skin graft transplanted onto female WT and *Tmem176a/b* DKO mice. Donor and recipient mice were matched for the presence or absence of *Tmem176a/b*. (**B**) Survival curves of WT and *Tmem176a/b* DKO mice injected with EG7, MCA101-sOVA or B16-OVA tumor cell line. (**C**) Schematic representation of EAE induction using MOG_35–55_ peptide or MOG_1–125_ protein. PT : Pertussis toxin. (**D**) EAE incidence and score (means±SEM) in WT and *Tmem176a/b* DKO mice using MOG_35–55_ peptide for immunization. (**E**) EAE incidence and score (means±SEM) in WT and *Tmem176a/b* DKO mice using MOG_1–125_ protein for immunization. (**F**) Schematic representation of DTH response using OVA_323–339_ peptide or whole OVA protein for immunization. (**G**) Left (control) and right footpad swelling (means±SEM) upon injection with the indicated heat-aggregated proteins in WT and *Tmem176a/b* DKO mice using OVA_323–339_ peptide for immunization. (**H**) Left (control) and right footpad swelling (means±SEM) upon injection with the indicated heat-aggregated proteins in WT and *Tmem176a/b* DKO mice using whole OVA protein for immunization.

To further test this hypothesis, we directly assessed antigen-specific T-cell proliferation in WT and DKO mice using naive OVA-specific CD8^+^ and CD4^+^ T cells (**Figure 5A**). It is important to note that cDCs are strictly required in this system to induce activation and proliferation of naive T cells^32^. As shown in **Figure 5B-C**, CD4^+^ T cell proliferation was markedly diminished in DKO mice in comparison to WT mice whereas CD8^+^ proliferation was not. Importantly, this alteration was not observed in *Tmem176a/b*^fl/fl^*Rorc*-Cre^+/–^ conditional mice (**Figure 5D**) but was replicated, although to a lesser extent than in germline DKO mice, in *Tmem176a/b*^fl/fl^*CD11c*-Cre^+/–^ mice (**Figure 5E**).

**Figure 5.**
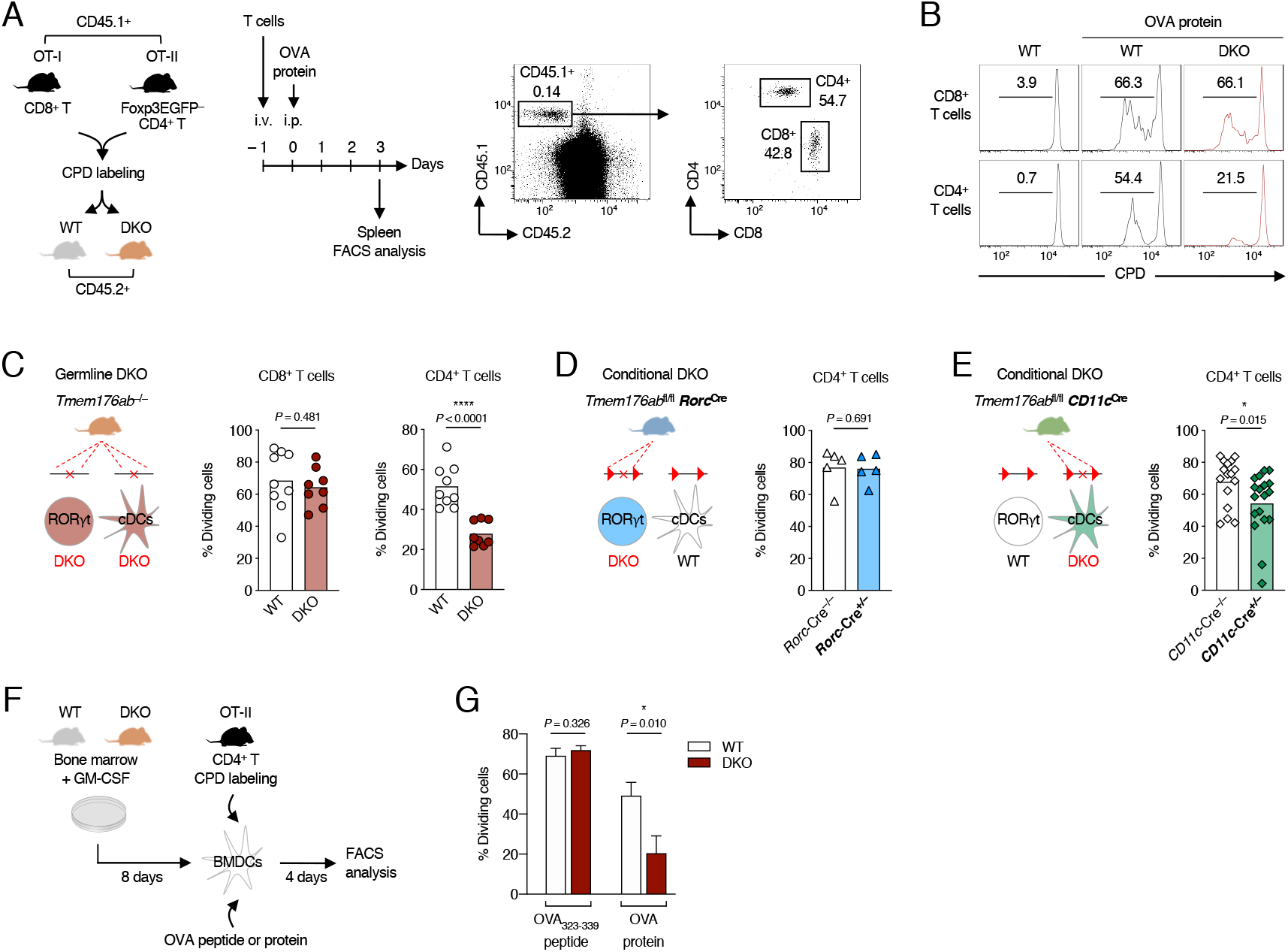
MHC I and II-mediated antigen presentation to T cells by *Tmem176a/b* DKO DCs. (**A**) Schematic representation of the in vivo antigen-specific T cell proliferation assay and gating strategy for tracking the injected OVA-specific T cells by flow cytometry. CPD : cell proliferation dye. (**B**) Representative histograms showing the level of proliferation (reflected by CPD dilution) of the OVA-specific CD8^+^ and CD4^+^ T cells following OVA protein injection in WT and *Tmem176a/b* DKO mice. (**C**) Quantification of OVA-specific CD8^+^ and CD4^+^ T cell proliferation in WT and *Tmem176a/b* DKO mice. (**D-E**) Quantification of OVA-specific CD4^+^ T cell proliferation in RORγt^+^ cell-restricted (**D**) or CD11c^+^ cell-restricted (**E**) conditional DKO mice. Bars indicate means and dots represent individual mice. Data were pooled from at least two independent experiments. (**F**) Schematic representation of the in vitro antigen-specific CD4^+^ T cell proliferation assay using BMDCs as antigen-presenting cells. (**G**) Quantification of CD4^+^ T cell proliferation following incubation of BMDCs from WT and *Tmem176a/b* DKO mice with OVA_323–339_ peptide or whole OVA protein. Data shown are means (±SD) of triplicates and are representative of two independent experiments.

In vitro antigen-specific CD4^+^ T-cell proliferation was also significantly decreased in the absence of *Tmem176a/b* using GM-CSF-induced bone-marrow derived DCs (BMDCs) only when OVA protein was used (**Figure 5F-G**), while surface expression of MHC II and co-stimulatory molecules remained unaltered (data not shown).

In conclusion, these results show that *Tmem176a/b* have an intrinsic function in DCs to allow efficient presentation of exogenous antigens onto MHC II molecules and priming of naive CD4^+^ T cells.

### Tmem176a/b-deficient cDCs exhibit normal antigen uptake and degradation but dysregulated H2-M expression selectively in cDC2

Antigen presentation by MHC II molecules is achieved through a series of complex events (depicted in **Figure 6A**) beginning with the uptake and mild degradation of exogenous antigens and including finely regulated processes in the specialized MHC II compartment (MIIC)^33^.

**Figure 6.**
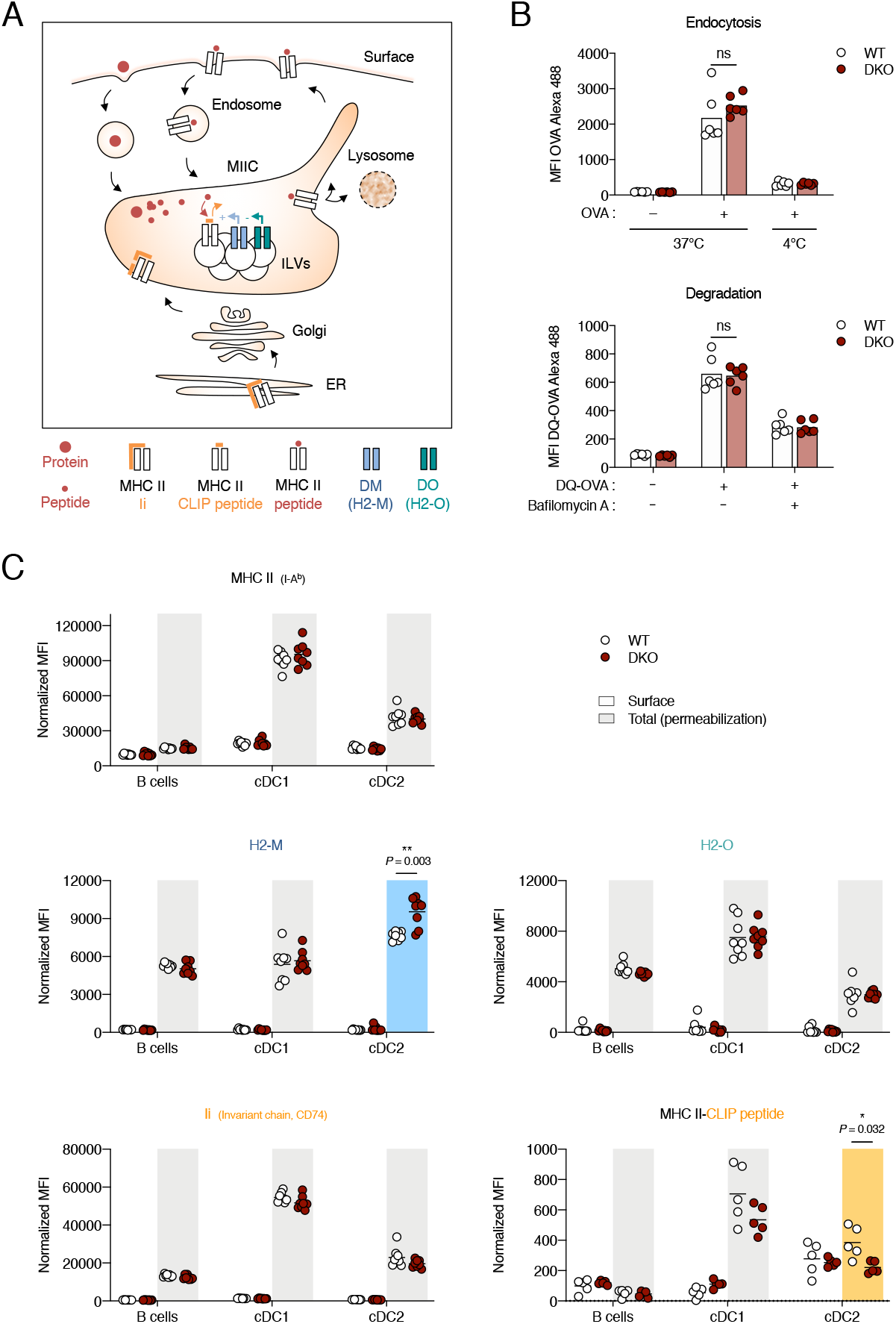
Antigen processing and expression of MHC II-associated molecules in *Tmem176a/b* DKO DCs. (**A**) Schematic overview of the MHC II pathway. The late endocytic compartment in which MHC II peptide loading occurs is referred to as the MHC II compartment (MIIC). Newly synthesized MHC II molecules in the endoplasmic reticulum (ER) bind to the invariant chain (Ii or CD74) that prevents premature loading with endogenous peptides before HLA-DM (H2-M in mouse) catalyzes the release of the class II-associated invariant chain peptide (CLIP) in the MIIC. Another MHC II-like protein, HLA-DO (H2-O in mouse), may add another level of regulation by inhibiting H2-M. ILVs : intraluminal vesicles. (**B**) In vitro antigen uptake and degradation by spleen cDCs from WT and *Tmem176a/b* DKO mice using Alexa 488-coupled OVA and DQ-OVA (that emits fluorescence upon degradation), respectively. Bars indicate means and dots represent individual mice. (**C**) Surface (white area) and intracellular (total, grey area) expression of MHC II, H2-M (αβ2 dimer), H2-O (β chain), Ii and MHC II-CLIP complex molecules in B cells, cDC1 and cDC2 populations from the spleen of WT and *Tmem176a/b* DKO mice. Bars indicate means and dots represent individual mice. Data were pooled from two independent experiments.

Both OVA endocytosis and degradation were similar in WT and DKO cDCs (**Figure 6B**), indicating that altered MHC II-mediated antigen presentation by *Tmem176a/b* DCs cannot be explained by a defect in the initial steps of antigen processing.

To determine whether *Tmem176a/b* deficiency could influence the surface or intracellular levels of key players in the MHC II pathway, we analyzed the expression of MHC II (I-A^b^), H2-M and H2-O in splenic B cells, cDC1 and cDC2 by flow cytometry (**Figure 6C**). As expected, cDC1 exhibited the highest levels of intracellular MHC II, Ii and H2-O whereas H2-M was primarily expressed in cDC2, an equilibrium concordant with the intrinsic efficiency of this subset in MHC II processing^30^. Although we did not detect aberrant expression of these molecules at the surface of DKO cells, we found that, intracellular H2-M was paradoxically and selectively over-expressed in the cDC2 subset of DKO mice compared to WT mice. Additionally, while Ii (invariant chain) expression was unaltered, the intracellular detection of the MHC II-CLIP peptide complex was not increased at the cell surface but was diminished intracellularly in cDC2 of DKO mice (**Figure 6C**, lower panels).

Taken together, these data suggest that *Tmem176a/b*-mediated function is directly involved in the MIIC for optimal MHC II antigen loading or trafficking.

### TMEM176A and TMEM176B traffic in dynamic vesicles between the Golgi apparatus and the endolysosomal compartments

To gain insight into the intracellular function of TMEM176A/B ion channels, we aimed to elucidate their subcellular localization that remains elusive as different studies reached different conclusions^5,6,34^. To this end, we used the Retention Using Selective Hooks (RUSH) system^35^, a two-state assay allowing fluorescence-based analysis of intracellular trafficking in living cells at physiological temperature (**Figure 7A)**. We performed dual-color imaging using multiple organelle-specific proteins or probes (**Figure 7B**) to track the intracellular fate of TMEM176A/B from the endoplasmic reticulum (ER). We used HeLa cells as they allow higher resolution of intracellular compartments compared to immune cells. Addition of biotin triggered a rapid change in the TMEM176B signal from a network of tubular elements characteristic of the ER to a pattern reminiscent of the Golgi apparatus but which rapidly evolved into multiple dynamic vesicles (**Figure 7C** and **Supplemental Video 1**). TMEM176A and TMEM176B exhibited a very similar intracellular dynamic, as measured by strong colocalization throughout the time of acquisition (**Figure 7D** and **Supplemental Video 2**).

**Figure 7.**
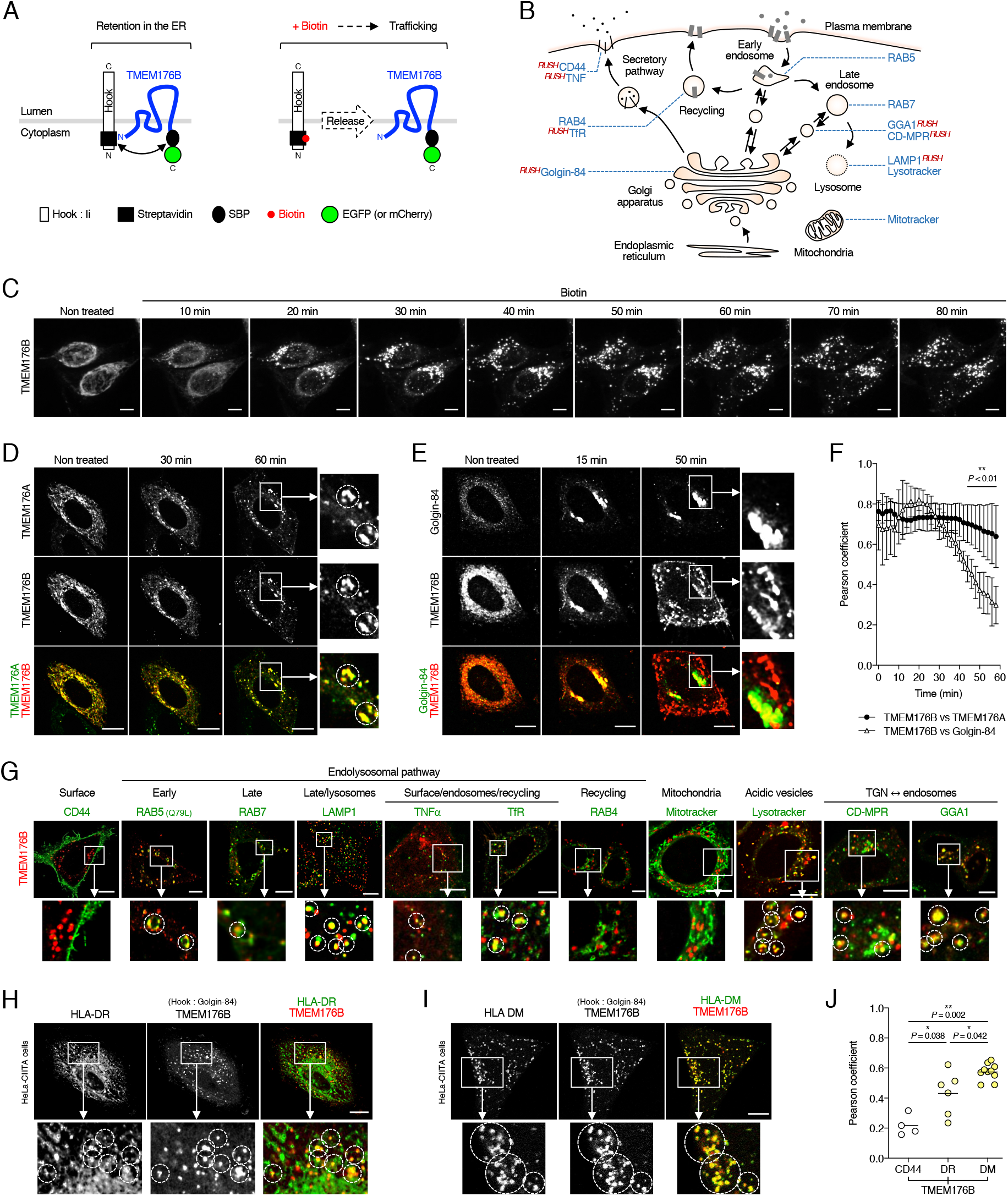
Intracellular trafficking of TMEM176A and TMEM176B using the RUSH system. (**A**) Principles of the retention using selective hooks (RUSH) system. In the setting we chose to implement for this two-state assay, the protein of interest is fused to the streptavidin binding protein (SBP) and is retained in the donor compartment (here the endoplasmic reticulum, ER) in which the hook (here an isoform of the invariant chain fused to the core streptavidin) remains localized. Synchronous release of the protein of interest is induced by addition of biotin and intracellular trafficking can be monitored by measuring fluorescent tags such as enhanced (E)GFP or mCherry signal by time-lapse confocal microscopy. (**B**) Schematic representation of the different intracellular compartments and associated markers analyzed. Proteins analyzed using RUSH constructs are indicated (**C**) Micrographs of HeLa cells expressing the TMEM176B-EGFP RUSH construct prior and after addition of biotin. Bar, 10 μm. (**D**) Dual-color analysis using the TMEM176A-EGFP and TMEM176B-mCherry RUSH constructs. Insets show higher magnifications of regions of interest. Circles show examples of colocalized signals. (**E**) Dual-color analysis using the TMEM176B-mCherry and Golgin-84-EGFP RUSH constructs. (**F**) Pearson’s correlation coefficients comparing TMEM176B with TMEM176A and Golgin-84 signals. (**G**) Dual-color analysis using the TMEM176B-mCherry or TMEM176B-EGFP RUSH constructs and the indicated genes or probes, >40 min after biotin addition. (**H-I**) Dual-color analysis in Hela-CIITA cells using the TMEM176B-mCherry RUSH construct with the YFP-associated HLA-DR (**H**) or HLA-DM (**I**) plasmids, >40 min after biotin addition. To avoid interference of the ER-resident mutated form of Ii (in the TMEM176B RUSH construct) on normal dynamic of the MHC II pathway, Ii was replaced by Golgin-84 as a hook in a new construct. To compensate for the low transfection efficiency of the DR/DM constructs, YFP^+^ cells were FACS-sorting 24 h before imaging. (**J**) Pearson’s correlation coefficients comparing TMEM176B with CD44, HLA-DR and HLA-DM signals. Each dot represents an individual cell.

Confirming our previous hypothesis that TMEM176B traffics through but does not accumulate in the Golgi apparatus^6^, it clearly separated from Golgin-84, a Golgi-resident protein. After 10–15 min of incubation with biotin, Golgin-84 reached the Golgi apparatus where it stayed after longer incubation time in contrast to TMEM176B (**Figure 7E-F** and **Supplemental Video 3**). We then examined a variety of markers depicted in **Figure 7G**. We did not, or only marginally, observe accumulation at the plasma membrane, a result that was confirmed by co-imaging with CD44. Interestingly, TMEM176B-bearing vesicles could interact with RAB5^+^ early endosomes as well as with RAB7^+^ late endosomes. We generated a LAMP1 (CD107a) RUSH construct to best reveal endolysosomes and also observed a strong association with TMEM176B trafficking during the post-Golgi time frame. Of interest, TMEM176B could be found colocalized with TNF or TfR (transferrin receptor), during the late events of endocytosis/recycling of these proteins. However, TMEM176B was not associated with RAB4, a marker of recycling endosomes. Although we cannot rule out alternate recycling pathways, these result suggest that TMEM176A/B follow a relatively selective route among the various vesicular compartments of the cell. In this line, TMEM176B did not traffic through mitochondria labeled by the Mitotracker probe but was significantly associated with Lysotracker that preferentially marks acidic vesicles. Moreover, we found some colocalization signals using a cation-dependent mannose-6-phosphate receptor (CD-MPR) RUSH construct but even stronger association with the monomeric clathrin adaptor GGA1, known to decorate the carrier vesicles budding from the TGN and merging toward the endosomes^36^.

Taken together, these data show that TMEM176A/B ion channels preferentially localize in the late endosomal compartment and in vesicular vesicles between the Golgi and the endolysosomal system.

### TMEM176B colocalizes with HLA-DM in MHC II-expressing cells

Given the requirement of *Tmem176a/b* in MHC II antigen presentation, the selective alteration of H2-M expression in cDC2 and the preferential trafficking of TMEM176A/B in the late endosomal compartment, we asked whether these ion channels could localize in the MIIC. To recapitulate the MHC II pathway in HeLa cells, we used the HeLa-CIITA cell line that stably expresses the transactivator CIITA^37^. To reveal MHC II (HLA-DR) and HLA-DM localization in these cells, we used plasmid constructs expressing YFP tagged to the beta or alpha chains of each molecule, respectively, as described in Zwarts *et al.*^38^. Colocalized signals were detected between TMEM176B and HLA-DR, mostly in intracellular vesicles (**Figure 7H**). However, a more pronounced association was observed with HLA-DM in intracellular compartments presumably highlighting the MIIC.

Thus, these results strongly suggest that TMEM176A/B exert their function directly in the MIIC to contribute to efficient MHC II peptide loading and/or trafficking.

## Discussion

Finely tuned ion influx and efflux result from various intricate interplays of multiple channels likely tailored for each type of cell and maturation status. The intriguing high expression of TMEM176A and TMEM176B cation channels in both RORγt^+^ lymphoid cells and DCs logically raises the question of their specific role in these two very distinct immune cell types. Based on expression and functional data, we reasoned that each gene has the potential to compensate for each other and that simultaneous targeting would be a requisite to avoid such redundancy.

We present here the first functional study of *Tmem176a/b* double KO (DKO) mice, either germline or conditional. Given the broad tissue expression of *Tmem176a/b*, the “floxed” conditional mouse represents an invaluable tool to achieve Cre-mediated cell-specific deletion and document the role of these ion channels in virtually any tissue or cell of interest. Indeed, there is a growing interest in understanding the role of these homolog genes that are over-expressed in a wide range of cell types other than immune cells including fibroblast subsets^39^, neurons^40^, adipocytes^41^ or tumor cells^42–44^, likely adapting a universal mechanism of intracellular ion flux regulation to their specific needs. Importantly, these *Tmem176a/b* DKO mice were generated directly in a pure genetic background (C57BL/6N) thus avoiding incorrect interpretations resulting from carryover of gene variants of a different background surrounding the targeted locus^45,46^.

The striking expression of *Tmem176a/b* in all type 3 immune cells and their dependency on the master transcription factor RORγt^15^ make these homologs promising candidates to uncover novel aspects of RORγt^+^ cell biology beyond their cytokine production and could represent a novel therapeutic entry point for treating immune-mediated diseases. However, our results indicate that RORγt-dependent intestinal repair and host defense functions are not compromised in the absence of *Tmem176a/b*. Furthermore, the normal development MOG_35–55_ peptide-induced EAE suggest that these genes are not required for the pathogenicity of Th17 in this model. Investigating the transcriptomic and epigenetic profiles of purified DKO ILC3s or Th17 cells may expose compensatory mechanisms notably involving the regulation of other ion channels that could sufficiently counterbalance the absence of *Tmem176a/b*. It is also tempting to speculate that these homologs could be required for IL-17/IL-22-independent functions in RORγt^+^ cells, including the regulation of anti-commensal effector CD4^+^ T-cells by ILC3s through MHC II-mediated inhibitory presentation^47,48^. Although *Tmem176a/b*^fl/fl^*Rorc*-Cre^+/–^ conditional mice did not exhibit increased CD4^+^ T cell activation and proliferation nor neutrophil accumulation in the colonic lamina propria in comparison to control mice (data not shown), the fact that ILC3s selectively share with DCs the expression of MHC II molecules is in favor of a pivotal role of *Tmem176a/b* in this adaptive function.

In view of the recent study by Segovia *et al.*^31^ using *Tmem176b* single KO mice or an ion channel inhibitory molecule, our results do not support the hypothesis that inhibiting TMEM176A/B-mediated ion flux could enhance anti-tumor CD8^+^ T-cell response in vivo. Moreover, we did not observe increased IL-1β production by *Tmem176a/b*-deficient BMDCs (data not shown). Although different experimental conditions could explain this discrepancy, one can speculate that TMEM176B must be targeted alone, leaving TMEM176A function intact, to obtain such phenotype, and that the effect of the inhibitory molecule is therefore achieved in a selective manner. Alternatively, because of the 129 genetic background origin of the *Tmem176b* single KO mouse^5^, confounding genetic factors could be invoked, despite >10 backcrosses onto the C57BL/6 background and the use of littermates^49^.

We initially reported that *Tmem176a/b* were highly expressed in cDCs but not in pDCs^10,17^, likely a consequence of E2-2-mediated repression as revealed by Ghosh *et al.* for *Tmem176a*^50^. Remarkably, recent mouse and human single-cell RNA-seq analysis highlighted these homologs as markers of selective DC subsets, both in mouse and human^51–53^. Notably, the association of *TMEM176B* expression with a subset of cDC2 in Binnewies *et al.*^53^ is concordant with our data in the mouse showing that *Tmem176a/b* are markedly over-expressed in cDC2 compared to cDC1. cDC2 exhibit an overall dominance in MHC II presentation in vivo resulting from the combination of their intrinsic efficiency^30,54^ and their favorable position within lymphoid tissues for antigen uptake^55^. Consistently, we found that *Tmem176a/b* deficiency selectively affected the capacity of DCs to prime naive CD4^+^ T cells but not CD8^+^ T cells in vivo. Our results point to a defect in the intracellular processing events for exogenous antigen presentation to MHC II molecules. However, this functional alteration is not complete and it is possible that, in the same manner as in ILC3s, compensatory mechanisms develop in DCs in the absence of *Tmem176a/b*. In support of this hypothesis, the fact that H2-M was found selectively over-expressed in cDC2 of DKO mice suggests an adaptation to alleviate a defect in the MIIC for peptide loading onto MHC II molecules. Alternatively, this expression may also reflect an incorrect intracellular localization of H2-M, therefore disrupting the optimal processes leading to MHC II presentation.

The analysis of TMEM176A/B intracellular dynamics enabled us to clearly delineate that they preferentially traffic in the late endolysosomal system in close relationship with the Golgi apparatus. In contrast with the limitations in sensitivity and the non-dynamic nature of classical immunostaining, the RUSH system was instrumental in revealing the route taken by TMEM176A/B from the ER and beyond the Golgi apparatus. However, because we focused on the first hour of trafficking after release in most of our analyses, we cannot exclude that TMEM176A/B can eventually reach other compartments over time.

The strong colocalization found with HLA-DM in HeLa-CIITA cells supports the hypothesis of a direct role in the MIIC. TMEM176A/B-mediated cation (Na^+^) efflux could participate in the regulated acidification of this compartment as a counterion conductance^56^. TMEM176A/B could be located on the limiting membrane of the MIIC or on intraluminal vesicles (ILVs) where a direct interaction with HLA-DM would be possible. In this regard, independently of their ion channel function, it is conceivable that these four-span transmembrane proteins act similarly to tetraspanin molecules to stabilize DM-MHC II interaction^57^. On the same note, given the reported genetic association between *TMEM176A* and HDL cholesterol levels in human^58^, TMEM176A/B function may be connected to cholesterol-containing microdomains for efficient MHC II trafficking^59^. High resolution imaging, FRET analysis or the characterization of organelle-specific disruption in DKO DCs could be informative to uncover the precise role of TMEM176A/B in the MHC II pathway.

In conclusion, while the intrinsic function of TMEM176A/B in RORγt^+^ cells remains to be further explored, we found that these cation channels play a substantial role in the MHC II pathway to ensure optimal naive CD4^+^ T cell priming by DCs. Remarkably, a recent study by the Rudensky group identified these genes by single-cell RNA sequencing as markers of the T-bet^−^ cDC2B subset both in mouse and human^22^, a finding that reinforces the hypothesis of a predominant role of TMEM176A/B in cDC2s. Together, these results also suggest that the generation of a *Tmem176a/b* reporter mouse could represent an invaluable tool to study cDC2 subsets in vivo.

## Materials and Methods

### Mice

*Tmem176a/b* double KO (DKO) mice were generated by a dual targeting approach using the CRISPR– Cas9 system as previously described^20^. DKO mice used in this study were generated in the C57BL/6N genetic background. Three consecutive backcrosses with C57BL/6N mice were performed before intercrossing heterozygous mice. To control for cage-dependent microbiota variations, WT and DKO mice were systematically co-housed directly after weaning following sex- and age-matching.

Conditional DKO mouse carrying a « floxed » *Tmem176a/b* allele (*Tmem176a/b*^fl^) were generated at Mouse Clinical Institute (Illkirch, France). Briefly, two consecutive rounds of ES cell (C57BL/6N genetic background) modifications using two independent selection cassettes were realized to insert LoxP sites on both sides of *Tmem176a* and *Tmem176b* first coding exons. F0 mouse chimera were crossed to a FlpO deleter mouse^60^ (pure C57BL/6N background) to remove the FRT- and F3-flanked Neomycin and Hygromycin selection cassettes abutted to the two LoxP sites. Allele transmission was verified on F1 mice before rederivation and housing in a SPF mouse facility. *Tmem176a/b*^fl/wt^ heterozygous mice were crossed to BAC transgenic *Rorc*(γt)-Cre mice (generated by Gérard Eberl^61^ and provided by Bernhard Ryffel) or *CD11c*-Cre mice (*Itgax*-Cre, generated by Boris Reizis^62^ and provided by Véronique Godot). Following intercrossing, co-housed, sex- and age-matched *Tmem176a/b*^fl/fl^ homozygous littermates carrying or not a transgenic Cre allele were used for experiments.

OT-I.*Ly5.1* homozygous mice were obtained by intercrossing OVA-specific TCR-transgenic OT-I mice (C57BL/6-Tg(TcraTcrb)1100Mjb/Crl) (Charles River) with *Ly5.1* mice (B6.SJL*-PtprcaPepcb*/BoyCrl) (Charles River).

OT-II.*Ly5.1.Foxp3EGFP* homozygous mice were obtained by intercrossing OVA-specific TCR-transgenic OT-II mice (C57BL/6-Tg(TcraTcrb)425Cbn/Crl) (Charles River) with *Ly5.1* mice (B6.SJL-*PtprcaPepcb*/BoyCrl) (Charles River) and *Foxp3EGFP* reporter mice (generated by Bernard Malissen^63^).

All mice used for experiments were between 8 and 25 weeks of age and kept under specific pathogen-free conditions. Experimental procedures were carried out in strict accordance with the protocols approved by the Commitee on the Ethics of Animal Experiments of Pays de la Loire and authorized by the French Government’s Ministry of Higher Education and Research.

### Chemically-induced acute colitis

Mice were given 2% dextran sulfate sodium (DSS) (36,000–50,000 MW, MP Biomedical) in drinking water ad libitum for 7 days followed by a recovery period without DSS. Mice were monitored and weighed daily.

### *Citrobacter rodentium* infection

*Citrobacter rodentium* (DBS100, ATCC 51459) were culture aerobically at 37°C overnight at 200 rpm in Luria-Bertani (LB) broth medium (MP Biomedicals) and then diluted 1:10 in fresh LB medium until the concentration of bacteria reached optical density 600. Mice were pre-treated with 750 mg/L metronidazole (Sigma) in 2.5% sucrose drinking water for 4 days as previously described^64^, followed by 3 days with regular drinking water. Mice were then fasted 8 h before the infection by oral gavage with 2×10^9^ colony-forming units (CFUs) of *C. rodentium* resuspended in sterile 0.9% NaCl. Bacterial concentration was assessed via serial dilution on LB agar plates to confirm the CFUs administered. Mice were monitored and weighed daily. Faeces were collected at days 0 and 6 post-infection for detection of *C. rodentium* by qPCR.

### Skin transplantation

Mice were anesthetized with a mixture of 5% xylazine (Rompun) and 18% ketamine in PBS (170 μL) injected intraperitoneally (8.5 mg/kg of xylazine and 76.5 mg/kg of ketamine per mouse). Square skin grafts (1 cm^2^) were prepared from the tail of male donors and transplanted on the shaved left flank of female recipients. The grafts were fixed to the graft bed with 10-12 interrupted sutures and were covered with protective tape. Mice were monitored every other day and graft rejection was defined as complete sloughing or a dry scab.

### Tumor growth models

EG7, MCA101-sOVA^65^ (provided by Clotilde Théry) and B16-OVA tumor cells were recovered from log phase in vitro growth and 1×10^6^ cells were injected subcutaneously in 50 μL of cold PBS into the flank skin of recipient mice. Tumor growth was measured in a blind fashion with a caliper and expressed as the area based on two perpendicular diameters. Mice were monitored daily and were euthanized when tumor size reached 289 mm^2^.

### Experimental Autoimmune-Encephalomyelitis (EAE)

For EAE induced with MOG peptide, mice were immunized subcutaneously at the base of the tail and lower flanks with 200 μg of MOG_35–55_ peptide (MEVGWYRSPFSRVVHLYRNGK, GenScript) emulsified in complete Freund’s adjuvant (Sigma) supplemented with Mycobacterium tuberculosis H37Ra (Difco Laboratories) at 8 mg/mL (400 μg/mL per mouse). Pertussis toxin (200 ng, Calbiochem) was injected intraperitoneally on the day of immunization and 2 days later.

For EAE induced with MOG protein^66^, mice were immunized subcutaneously at the base of the tail and lower flanks with 500 μg of mMOGTag protein (mouse MOG_1–125_ extracellular domain fused to a tag for stability and purification purposes) provided by Steven Kerfoot and emulsified in complete Freund’s adjuvant. Pertussis toxin (250 ng) was injected intraperitoneally on the day of immunization and 2 days later.

Mice were scored daily for EAE clinical signs on a scale of 0–5 : 0, no disease; 1, complete limp tail; 2, limp tail with unilateral hindlimb paralysis; 3, bilateral hindlimb paralysis; 4, bilateral hindlimb paralysis and forelimb weakness (end point). The observer was blinded to the genotype during the scoring.

### Delayed-type hypersensitivity (DTH) assay

Mice were immunized subcutaneously at the base of the tail and lower flanks with 50 μg of whole OVA protein (grade V, Sigma) or OVA_323–339_ (ISQAVHAAHAEINEAGR) peptide (GenScript) emulsified in complete Freund’s adjuvant (Sigma). After 7 days, mice were challenged with 250 μg of heat-aggregated OVA (2 min incubation at 100°C) injected (20 μL, s.c.) in the right hind footpad whereas the left hind footpad received 250 μg of heat-aggregated BSA (Sigma) as a control for non-specific inflammation. Footpad thickness was measured prior to, 24 h and 48 h after injection with an electronic digital micrometer. The observer was blinded to the genotype during the scoring.

### Flow cytometry analysis and cell sorting

Antibodies and panels used in this study for FACS analysis and cell sorting are listed in Supplemental Table 1. Red blood cells were lysed with ammonium chloride. Small intestine and colon lamina propria (siLP an cLP) cells were prepared as previously described^6^. Before all stainings, dead cells were marked for exclusion using Fixable Viability Dye eFluor 506 (eBioscience) or DAPI (Thermo Fisher Scientific) followed by Fc receptor blocking using CD16/32 antibody (BD Biosciences). Intracellular stainings were realized using eBioscience Foxp3 / Transcription Factor Staining Buffer Set except for MHC II-CLIP peptide staining where cells were fixed using 4% PFA before permeabilization and staining using a 0.1% saponine, 1% BSA solution in PBS. FACS analyses were performed using BD FACS Canto II or a BD LSRFORTESSA X-20 (BD Biosciences) and FlowJo (Treestar) software. For mean fluorescence intensity (MFI) analysis, values were ajusted by substracting the basal signal from fluorescence minus one (FMO) staining for each marker. Absolute cell numbers were determined using CytoCount microspheres (Dako, Agilent). Total and surface expression analysis of MHC II, H2-M, H2-O, Ii (invariant chain, CD74) and MHC II-CLIP peptide complex in spleen cDC1 and cDC2 were performed with or without a fixation/permeabilization step, respectively, and by gating on B cells (B220^+^CD11c^−^), cDC1 (B220^−^CD11c^+^CD11b^−^CD8α^+^) or cDC2 (B220^−^CD11c^+^CD11b^+^CD8α^−^). H2-M (αβ2 dimer) staining was revealed using a FITC-conjugated anti-rat IgG1 antibody. Alexa Fluor 647-conjugated anti-H2-Oβ was provided by Liza Denzin^67^.

Cells were FACS-sorted using BD FACSAria II (BD Biosciences). For verification of Cre-induced loxP recombination, CD11b^−^TCRb^+^CD4^+^CD8^−^ T cells were FACS-sorted from the lamina propria of the small intestine of mice treated as previously described^21^ with 20 μg anti-CD3 (145-2C11, kindly provided by J.A. Bluestone), i.p., at days –3 and –1 before sacrifice. CD11c^high^MHC II^+^ cDCs were FACS-sorted at high purity (>98%) from the spleen of naive mice after enrichment of CD11c^+^ cells using a PE-conjugated anti CD11c antibody and magnetic-activated cell sorting (Anti-PE MicroBeads, Miltenyi Biotec). For in vitro functional analysis, intestinal CD3^+^CD5^+^CD4^+^ T cells and CD3^−^CD5^−^ CD127^+^ ILCs (first gated on CD45^+/low^CD11b/c^−^CD19^−^CD90^+^ cells) were FACS-sorted from the lamina propria of the small intestine and colon (pooled). For spleen cDCs used in epigenetic analysis and in vitro culture, CD11c^high^MHC II^+^ cells were purifed as described above. For RT-qPCR analysis, B cells (CD19^+^B220^+^CD11c^−^), pDCs (CD19^−^B220^+^CD11c^+^) cDC1 (CD19^−^B220^−^CD11c^high^CD11b^−^CD8α^+^) and cDC2 (CD19^−^B220^−^CD11c^high^CD11b^+^CD8α^−^) were FACS-sorted from the spleen of WT mice. OVA-specific CD8^+^ T cells were purified from the spleen of OT-I.*Ly5.1* mice using CD8a^+^ T Cell Isolation Kit II (Miltenyi Biotec). OVA-specific CD4^+^EGFP^−^ cells T conventional cells were FACS-sorted from the spleen of OT-II.*Ly5.1.Foxp3EGFP* mice after enrichment using CD4^+^ T Cell Isolation Kit (Miltenyi Biotec).

### In vivo antigen-specific T-cell proliferation assay

OVA-specific naive CD8^+^ T and CD4^+^ T conventional cells from OT-I.*Ly5.1* and OT-II.*Ly5.1.Foxp3EGFP* mice, respectively, were labeled with Cell Proliferation Dye (CPD) eFluor 670 (eBioscience) and co-injected (i.v.) at a 1:1 ratio (total of 1–2×10^6^ cells per mouse). One day later, recipient mice were administered (i.p.) 100 μg EndoFit Ovalbumin protein (InvivoGen). After three days, spleens were harvested and proliferation (CPD dilution) of injected cells (CD45.1^+^CD8^+^ or CD45.1^+^CD4^+^) was assessed by flow cytometry.

### BMDC generation and in vitro antigen-specific T-cell proliferation assay

Bone marrow cells were cultured (0.5×10^6^ cells per mL) in the presence of 20 ng/mL GM-CSF (Miltenyi Biotec) in complete RPMI medium. By day 8, the cells, referred to as BMDCs, were harvested and incubated in 96-well plate (1×10^4^ cells per well) with 250 μg/mL EndoFit Ovalbumin protein (InvivoGen) or 10 μg/mL OVA_323–339_ (ISQAVHAAHAEINEAGR) peptide (GenScript). After 5 h, the cells were washed three times and CPD-labeled CD4^+^ Foxp3^−^ T cells purified from OT-II.*Ly5.1.Foxp3EGFP* mice were added and proliferation (CPD dilution) was assessed by flow cytometry 3 days later.

### In vitro stimulation and analysis of cDCs and BMDCs

Purified spleen cDCs were plated in 96-well plate at 1×10^5^ cells and incubated with or without 0.5 μg/mL LPS (Sigma) for 16 h before the quantification of IL-12p40 and IL-6 in the supernatant by ELISA (BD Biosciences). For phenotypic analysis (MHC II, MHC I, CD80, CD86) by flow cytometry, bulk spleen cells (comprising cDC1 and cDC2 identified using the markers CD11c, CD11b and CD8α as indicated) or BMDCs were stained freshly of stimulated for 6 h with 0.5 μg/mL LPS.

### In vitro stimulation of CD4^+^ T cells and ILCs

Purified intestinal CD4^+^ T cells and ILCs were cultured in vitro in 96-well plate (10,000 cells per well in triplicates) in complete medium (Gibco, Thermo Fisher Scientific) for 18 hours in the presence of anti-CD3/CD28 Dynabeads (Thermo Fisher Scientific) at a ratio of 2:1 or IL-23 ± IL-7, IL-2, IL-1β (R&D Systems) at 50 ng/mL (except IL-2 at 50 IU/mL), respectively. IL-17A, IL-17F, IL-22 (R&D Systems) and IFNγ (BD Biosciences) were then measured in the supernatant by ELISA.

### In vitro antigen uptake and degradation assays

Antigen endocytosis and degradation in cDCs (CD11c^+^MHCII^+^) were assessed by flow cytometry after incubating one million bulk splenocytes in 96-well plate with 50 μg/mL OVA-FITC or DQ-OVA (Invitrogen, Thermofisher Scientific), respectively. As controls, cells were incubated at 4°C or treated with Bafilomycin A (Sigma).

### ChIP-seq experiments

Highly purified spleen cDCs (1×10^6^ cells from two pooled mice for each preparation) were resuspended in 40 μL of PBS. Cells were lysed and chromatin was fragmented with 300 units of Micrococcal nuclease (MNase) (M0247S, New England Biolabs) per well for 10 min at 37°C. After full speed centrifugation, supernatants were collected and filled up to 400 μL. Two μg of anti-H3K27ac (39133, Active Motif) was used for immunoprecipitation overnight at 4°C. Twenty five μL of G-protein dynabeads (Invitrogen) were added for rotation for 4 h at 4°C. Beads were then washed twice with 200 μL wash buffers with increasing salt concentration. ChIP beads were eluted in 50 μL of ChIP elution buffer (50 mM Tris-HCl pH 7.5, 10 mM EDTA, 1% SDS). ChIP and input samples were digested with 250 μg/mL proteinase K (GEXPRK006R, Eurobio) in 50 μL TE buffer for 1 h at 63°C. ChIP DNA was purified using phenol chlorophorm. Libraries were then prepared according as previously described^68^. Libraries were verified and equimolar pools were sequenced on a NextSeq 500 (75 bp single-end).

### ChIP-seq analysis

Single-end reads were mapped to the mm10 genome by the BWA algorithm and reads mapping to non-canonical and mitochondrial chromosomes were also removed. For each sample, ChIP-seq peaks were detected using DFilter at a *P*-value threshold of 1×10^−6^. All samples passed the quality controls (Fraction of reads in peaks [FRiP] > 3% and non-redundancy fraction [NRF] > 0.9). A set of consensus peaks was then obtained by taking the union of all peaks and counting reads these peaks using Bedtools. To perform Differential Peak Calling, differentially acetylated (DA) peaks were determined using edgeR after a counts per million (cpm) normalization. DA peaks were defined with a Benjamini-Hochberg Q-value ≤5%. For heatmap representation, peaks were rlog transformed. To determine gene ontology enrichment in up-regulated peaks the GREAT tool was used.

### Quantitative PCR

Quantitative PCR (qPCR) was performed using ViiA 7 Real-Time PCR System and Fast SYBR Green Master Mix reagent (Applied Biosystems, Thermo Fisher Scientific). Primer sequences are listed in Supplemental Table 1).

For *Citrobacter rodentium* bacterial load quantification, genomic DNA (gDNA) from faeces was purified using QIAamp DNA Stool Mini Kit (Qiagen) and the *EspB* gene was detected using specific primers and normalized with total bacterial gDNA (*16S* gene) using the 2^−ΔΔCt^ method.

For mRNA quantification, total RNA was isolated using RNeasy Mini Kit or Micro Kit (Qiagen). Reverse transcription was performed using M-MLV Reverse Transcriptase and random primers following manufacturer’s instructions (Thermo Fisher Scientific). Gene-specific primers were designed over different exons to prevent amplification of genomic DNA. Gene expression was normalized to glyceraldehyde 3-phosphate dehydrogenase (*Gapdh*) and expressed in arbitrary units using the 2^−ΔΔCt^ method.

### RUSH system

RUSH experiments were realized as previously described by the Perez group^35^. Plasmid constructs are listed in Supplemental Table 1. When indicated, 250 nM MitoTracker Deep Red or 100 nM LysoTracker Deep Red (Thermo Fisher Scientific) were added 30 min before imaging. HeLa cells (1.5×10^4^) were seeded into a μ-Slide 8 Well Chamber Slide (Ibidi) and transfected using Lipofectamine LTX Plus Reagent (Thermo Fisher Scientific). After 24 h, the medium was replaced by medium containing 40 μM of D-biotin (Sigma). The initial time-lapse acquisition characterizing TMEM176B was performed at Institut Curie (Paris, France) with a thermostat controlled chamber using an Eclipse 80i microscope (Nikon) equipped with a spinning disk confocal head (Perkin) and an Ultra897 iXon camera (Andor). Subsequent RUSH analyses were performed at MicroPICell facility (Nantes, France) with a Confocal Nikon A1 RS microscope. To analyze molecule colocalization, we used the plugin Colocalization Studio that contains pixel-based methods that were introduced in section Pixel-Based Methods in the Icy platform. HeLa-CIITA (generated by Philippe Benaroch^37^) were used for the analysis of HLA-DR and HLA-DM.

### Statistical analysis

Statistical analyses were performed using Graphpad Prism (La Jolla, CA, USA) using the Mann-Whitney test or, for survival rates, using the Log-rank (Mantel-cox) test. *P* values < 0.05 were considered significant.

## Acknowledgments

This work was supported by IHU-Cesti, Nantes Métropole and Région des Pays de la Loire, Paris Scientifiques Régionaux. This work was realized in the context of the Labex IGO program supported by the Agence Nationale de la Recherche (ANR-11-LABX-0016-01). This work was supported by the Fondation pour la Recherche Médicale, grant number ECO20160736078, to Melanie Lancien. We are grateful to Philippe Hulin, Steven Nedellec and Magalie Feyeux from the MicroPICell imagery core facility (Nantes, France) for excellent assistance with confocal microscopy and to Claire Usal, Pierre Pajot and Jean-Marc Merieau for mouse housing and experimental help.

Work performed in the F. Perez laboratory was funded by Centre National de la Recherche Scientifique, the Fondation pour la Recherche Medicale (FRM DEQ20120323723), the Labex CellTisPhyBio, and the Agence Nationale de la Recherche (ANR-12-BSV2-0003-01). The laboratory of F. Perez is part of Labex CelTisPhyBio (11-LBX-0038) and Idex Paris Sciences et Lettres (ANR-10-IDEX-0001-02 PSL). The authors acknowledge the Cell and Tissue Imaging Facility (PICT-IBiSA), Institut Curie, a member of the French National Research Infrastructure, France-BioImaging (ANR10-INBS-04).

## Author contributions

C.L., M.L. and M.C.C. designed and supervised the research. M.L., G.Bienvenu, L.G, S.S., E.M., S.R., A.Molle, C.F., G.Beriou P.K., S.A.R, V.D.S., G.Boncompain, J.P. and C.L. performed experiments and analyzed data. A.E., C.F., L.B-D and F.C., performed experiments. S.K., G.M. and F.P. contributed to key research tools and experiment design. A.Moreau., E.C., S.C., F.P., R.J. and J.P. helped with the study design and data interpretation. C.L., M.L., J.P., A.Molle and C.F. wrote the paper.

## List of Supplemental data

**Figure S1. *Tmem176a* and *Tmem176b* expression in the immune system.**

**Figure S2. Gating strategy for T cell and ILC FACS analysis and gene expression analysis by RT-qPCR analysis.**

**Figure S3. Gating strategy for bone-marrow progenitor analysis by flow cytometry**.

**Table S1. List of antibodies, oligonucleotides and plasmids used in this study.**

**Video S1. Real-time imaging of the synchronized trafficking of TMEM176B using the RUSH system (corresponds to Figure 7C).**

**Video S2. Dual color, real-time imaging of the synchronized trafficking of TMEM176B and TMEM176A using the RUSH system (corresponds to Figure 7D).**

**Video S3. Dual color, real-time imaging of the synchronized trafficking of TMEM176B and TMEM176A using the RUSH system (corresponds to Figure 7E).**

