## Supplemental Figures 1, 2, 3 and legends of Videos 1, 2, 3 for "Dendritic cells require TMEM176A/B ion channels for optimal MHC II antigen presentation to naive CD4^+^ T cells"

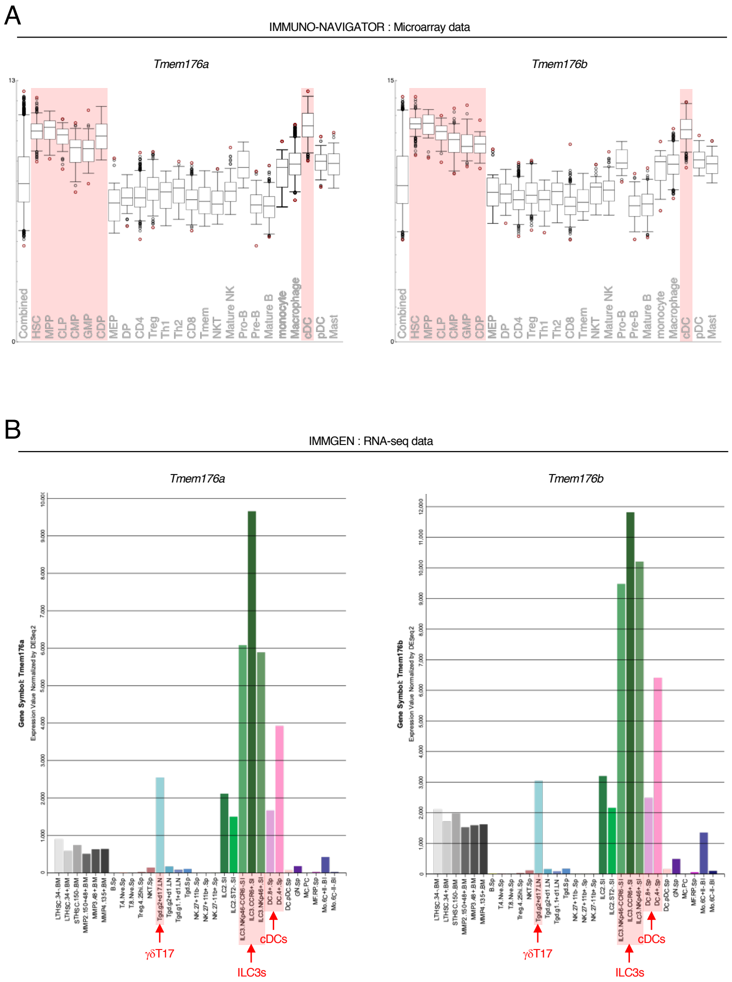


**Figure S1. *Tmem176a* and *Tmem176b* expression in the immune system.** mRNA expression in different panels of purified immune cells from (**A**) Immuno-Navigator microarray data (https://genomics.virus.kyoto-u.ac.jp/immuno-navigator/) and (**B**) Immgen RNA-seq data (http://www.immgen.org/). The populations exhibiting the highest relative expression are highlighted in red. HSC : hematopoietic stem cells, MPP : multipotent progenitors, CLP : common lymphoid progenitors, CMP : common myeloid progenitors, GMP : granulocyte-macrophage progenitors, CDP : common dendritic progenitors, MEP : megakaryocyte-erythroid progenitors, DP : Thymic double positive cells, CD4 : $CD4^+$ T cells, Th1 : T helper 1, Th2 : T helper 2, CD8 : $CD8^+$ T cells, Tmem : memory T cells, NKT : natural killer T cells, Mature NK : mature natural killer cells, cDC : conventional dendritic cells, pDC : plasmacytoid dendritic cells, Mast : Mastocytes.

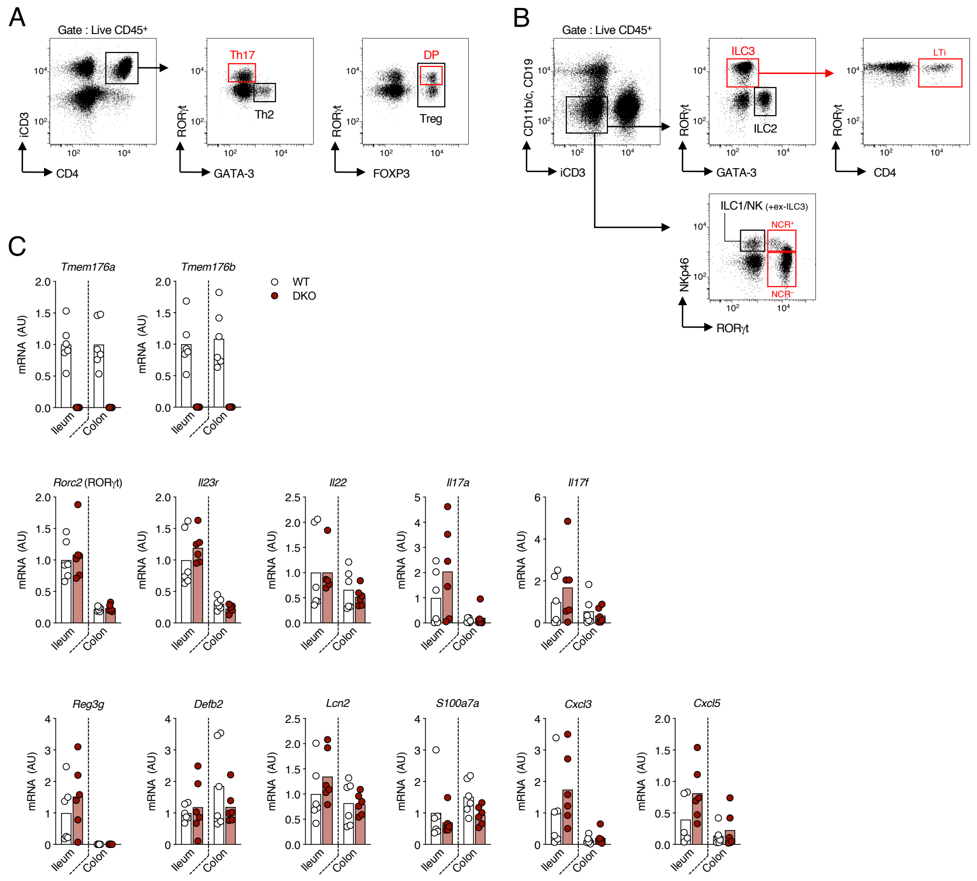


**Figure S2. Gating strategy for T cell and ILC FACS analysis and gene expression analysis by RT-qPCR analysis.** (**A-B**) Gating strategy for the analysis (shown in Figure 2A) of T (**A**) and ILC (**B**) subsets in the small intestine (siLP), colon (cLP) lamina propria and spleen by flow cytometry. Shown are examples of WT siLP analysis. iCD3 indicates that the anti-CD3ε staining was performed intracellularly along with the transcription factor stainings. DP : double positive cells, NCR : natural-cytotoxicity-receptor cells, LTi : lymphoid-tissue-inducer cells. (**C**) RT-qPCR analysis of *Tmem176a*, *Tmem176b*, RORγt-related genes and IL-22/IL-17 target genes in the ileum and colon of *Tmem176a/b* DKO mice. Pieces of ileum and colon were collected from WT and *Tmem176a/b* DKO mice and total RNA was extracted for RT-qPCR relative expression analysis of the indicated genes. Bars indicate means and dots represent individual mice.

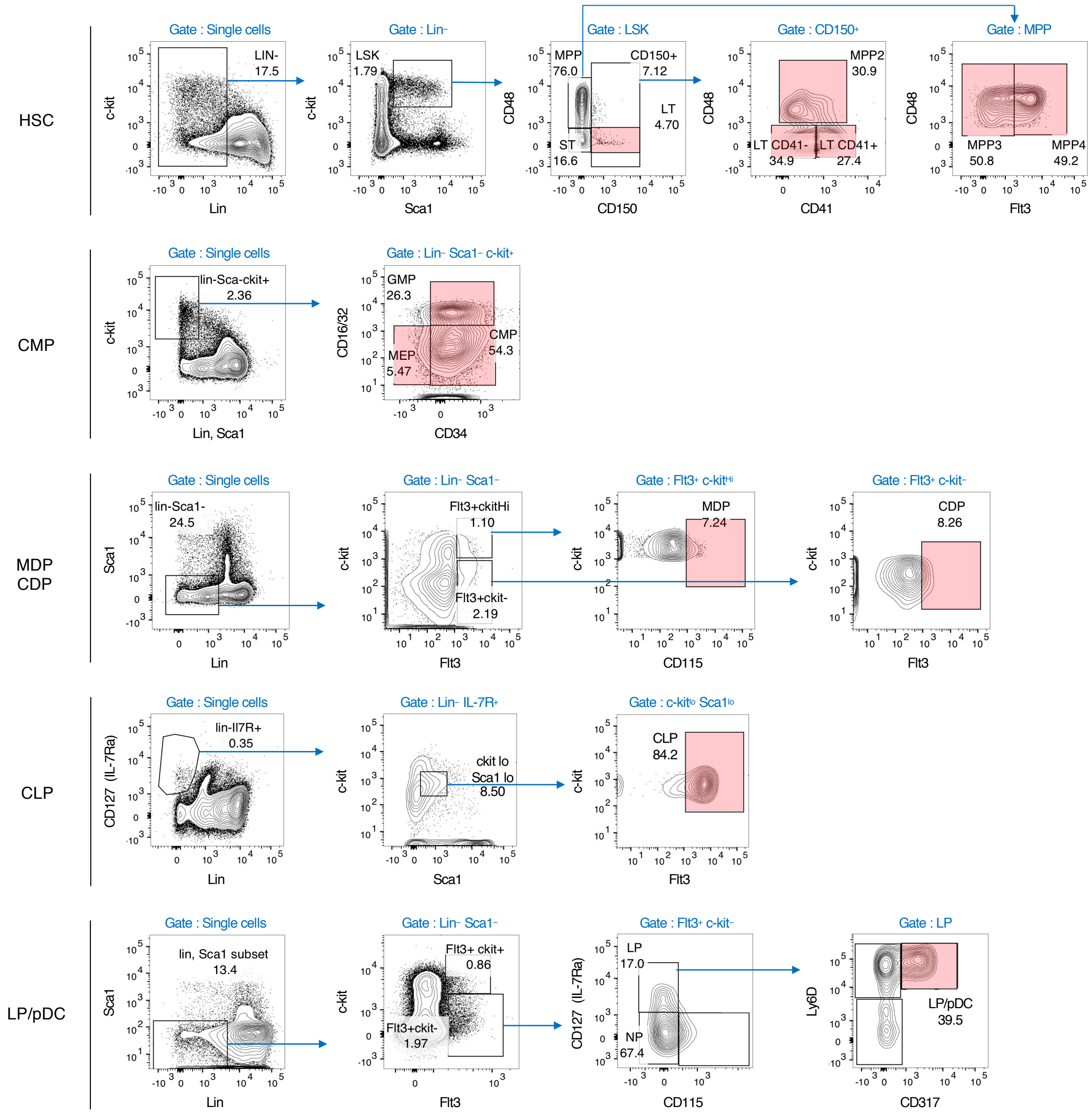


**Figure S3. Gating strategy for bone-marrow progenitor analysis by flow cytometry**. Gates highlighted in red indicate the progenitor populations (schematically represented in Figure 1B) that were quantified in WT and *Tmem176a/b* DKO mice (shown in Figure 3A).

**Table S1. List of antibodies, oligonucleotides and plasmids used in this study.**

**Video S1. Real-time imaging of the synchronized trafficking of TMEM176B using the RUSH system (corresponds to Figure 7C).** After 20 h of expression, at time 00:00, release of the reporter TMEM176B-SBP-EGFP was induced by addition of biotin and monitored using a spinning disk confocal microscope.

**Video S2. Dual color, real-time imaging of the synchronized trafficking of TMEM176B and TMEM176A using the RUSH system (corresponds to Figure 7D).** After 20 h of expression, at time 00:00, release of the reporters TMEM176B-SBP-mCherry (red) and TMEM176A-SBP-EGFP (green) was induced by addition of biotin and monitored using a confocal microscope.

**Video S3. Dual color, real-time imaging of the synchronized trafficking of TMEM176B and TMEM176A using the RUSH system (corresponds to Figure 7E).** After 20 h of expression, at time 00:00, release of the reporters TMEM176B-SBP-mCherry (red) and Golgin-84-SBP-EGFP (green) was induced by addition of biotin and monitored using a confocal microscope.
