## Supplemental Table 1 for "Dendritic cells require TMEM176A/B ion channels for optimal MHC II antigen presentation to naive CD4^+^ T cells"

### Lancien *et al.* - Table S1 List of antibodies, oligonucleotides and plasmids used in this study

#### Antibodies used in this study

(intra) indicates that the antibody was used following a fixation/permeabilisation step to allow intracellular staining

##### Figure 1E-F : Small intestine (si) $CD4^{+}$ T cell and spleen cDC FACS-sorting from conditional DKO mice

**si$CD4^{+}$ T cells**

| Specificity | Clone | Fluorochrome or biotin | Source |
|---|---|---|---|
| CD16/32 | 2.4G2 | - | BD Biosciences |
| TCRb | H57-597 | APC | eBioscience |
| CD11b | M1/70 | APC | BD Biosciences |
| CD4 | RM4-5 | PE-Cy7 | BD Biosciences |
| CD8a | 53-6.7 | FITC | BD Biosciences |

**Spleen cDCs**

| Specificity | Clone | Fluorochrome or biotin | Source |
|---|---|---|---|
| CD16/32 | 2.4G2 | - | BD Biosciences |
| CD11c | HL3 | PE | BD Biosciences |
| I-$A^{b}$ (MHC II) | AF6-120.1 | APC | eBioscience |

##### Figures 2A and S2A-B : Intestinal $CD4^{+}$ T cell and ILC analysis

**$CD4^{+}$ T cells**

| Specificity | Clone | Fluorochrome or biotin | Source |
|---|---|---|---|
| CD16/32 | 2.4G2 | - | BD Biosciences |
| CD45.2 | 104 | APC-Cy7 | BD Biosciences |
| CD4 | RM4-5 | V450 | BD Biosciences |
| CD8a | 53-6.7 | PE-Cy7 | BD Biosciences |
| CD3e (intra) | 145-2C11 | PerCP-Cy5.5 | BD Biosciences |
| RORgt (intra) | Q31-378 | Alexa Fluor 647 | BD Biosciences |
| GATA-3 (intra) | TWAJ | Alexa Fluor 488 | eBioscience |
| FOXP3 (intra) | FJK16S | PE | eBioscience |

**ILCs**

| Specificity | Clone | Fluorochrome or biotin | Source | |
|---|---|---|---|---|
| CD16/32 | 2.4G2 | - | BD Biosciences | |
| CD45.2 | 104 | APC-Cy7 | BD Biosciences | |
| CD11b | M1/70 | PE | BD Biosciences | Lineage |
| CD11c | HL3 | PE | BD Biosciences | |
| CD19 | 1D3 | PE | BD Biosciences | |
| CD3e (intra) | 145-2C11 | PerCP-Cy5.5 | BD Biosciences | |
| RORgt (intra) | Q31-378 | Alexa Fluor 647 | BD Biosciences | |
| GATA-3 (intra) | TWAJ | Alexa Fluor 488 | eBioscience | |
| CD4 | RM4-5 | PE-Cy7 | BD Biosciences | |
| NKp46 (CD335) | 29A1.4 | eFluor 450 | eBioscience | |

##### Figure 2B : Intestinal $CD4^{+}$ T cell and ILC FACS-sorting for in vitro assay

| Specificity | Clone | Fluorochrome or biotin | Source |
|---|---|---|---|
| CD16/32 | 2.4G2 | - | BD Biosciences |
| CD45.2 | 104 | APC-Cy7 | BD Biosciences |
| CD11b | M1/70 | PE | BD Biosciences |
| CD11c | HL3 | PE | BD Biosciences |
| CD19 | 1D3 | APC | BD Biosciences |
| CD90.2 | 53-2.1 | eFluor 450 | eBioscience |
| CD3e | 145-2C11 | FITC | BD Biosciences |
| CD5 | 53-7.3 | FITC | BD Biosciences |
| CD4 | RM4-5 | PerCP-Cy5.5 | BD Biosciences |
| CD127 | SB/199 | PE-Cy7 | BD Biosciences |

##### Figures 3A and S3 : Bone-marrow HSC and progenitor analysis

**HSC**

| Specificity | Clone | Fluorochrome or biotin | Source | |
|---|---|---|---|---|
| CD16/32 | 93 | - | BioLegend | |
| CD3 | 145-2C11 | Biotin | BD Biosciences | Lineage |
| CD4 | GK1.5 | Biotin | BD Biosciences | |
| CD11b | M1/70 | Biotin | BD Biosciences | |
| CD11c | N418 | Biotin | BioLegend | |
| CD19 | 6D5 | Biotin | BioLegend | |
| B220 | RA3-6B2 | Biotin | BD Biosciences | |
| Gr-1 | RB6-8C5 | Biotin | BD Biosciences | |
| CD49b | DX5 | Biotin | BioLegend | |
| Ter119 | Ter119 | RB6-8C5 | BD Biosciences | |
| CD8 | 53-6.7 | Biotin | BioLegend | |
| Streptavidin | - | eFluor 710 | eBioscience | |
| c-kit | 2B8 | APC | BioLegend | |
| Sca1 | D7 | PE-Cy7 | BD Biosciences | |
| CD48 | HM48-1 | FITC | BioLegend | |
| CD150 | TC15-12F12.2 | BV421 | BioLegend | |
| Flt3 | A2F10 | PE | eBioscience | |
| CD41 | MWReg30 | BV510 | BioLegend | |

**CMP/MEP/GMP**

| Specificity | Clone | Fluorochrome or biotin | Source | |
|---|---|---|---|---|
| CD3 | 145-2C11 | Biotin | BD Biosciences | |
| CD4 | GK1.5 | Biotin | BD Biosciences | |
| CD11b | M1/70 | Biotin | BD Biosciences | |
| CD11c | N418 | Biotin | BioLegend | |
| CD19 | 6D5 | Biotin | BioLegend | |
| B220 | RA3-6B2 | Biotin | BD Biosciences | Lineage |
| Gr-1 | RB6-8C5 | Biotin | BD Biosciences | |
| CD49b | DX5 | Biotin | BioLegend | |
| Ter119 | Ter119 | Biotin | BD Biosciences | |
| CD8 | 53-6.7 | Biotin | BioLegend | |
| Sca1 | D7 | Biotin | eBioscience | |
| Streptavidin | - | Alexa Fluor 405 | Life technologies | |
| CD34 | MEC14.7 | PE | BioLegend | |
| c-kit | 2B8 | APC | BioLegend | |
| CD16/32 | 93 | PE-Cy7 | BioLegend | |

**MDP/CDP/CLP**

| Specificity | Clone | Fluorochrome or biotin | Source | |
|---|---|---|---|---|
| CD16/32 | 93 | - | BioLegend | |
| CD3 | 145-2C11 | Biotin | BD Biosciences | |
| CD4 | GK1.5 | Biotin | BD Biosciences | |
| CD11b | M1/70 | Biotin | BD Biosciences | |
| CD11c | N418 | Biotin | BioLegend | |
| CD19 | 6D5 | Biotin | BioLegend | |
| B220 | RA3-6B2 | Biotin | BD Biosciences | Lineage |
| Gr-1 | RB6-8C5 | Biotin | BD Biosciences | |
| CD49b | DX5 | Biotin | BioLegend | |
| Ter119 | Ter119 | Biotin | BD Biosciences | |
| CD8 | 53-6.7 | Biotin | BioLegend | |
| Streptavidin | - | Alexa Fluor 405 | Life technologies | |
| Flt3 | A2F10 | PE | eBioscience | |
| c-kit | 2B8 | APC-Cy7 | BioLegend | |
| CD115 | AFS98 | APC | eBioscience | |
| CD127 | A7-R34 | PE-Dazzle594 | BioLegend | |
| Sca1 | D7 | PE-Cy7 | BD Biosciences | |

**LP/pDC**

| Specificity | Clone | Fluorochrome or biotin | Source | |
|---|---|---|---|---|
| CD16/32 | 93 | - | BioLegend | |
| CD3 | 145-2C11 | Biotin | BD Biosciences | |
| CD4 | GK1.5 | Biotin | BD Biosciences | |
| CD11b | M1/70 | Biotin | BD Biosciences | |
| CD11c | N418 | Biotin | BioLegend | |
| CD19 | 6D5 | Biotin | BioLegend | |
| B220 | RA3-6B2 | Biotin | BD Biosciences | |
| Ly6C (CD59) | ER-MP20 | Biotin | BMA Biomedicals | Lineage |
| Gr-1 | RB6-8C5 | Biotin | BD Biosciences | |
| NK1.1 | PK136 | Biotin | BioLegend | |
| Ter119 | Ter119 | Biotin | BD Biosciences | |
| CD8 | 53-6.7 | Biotin | BioLegend | |
| Streptavidin | - | eFluor 710 | eBioscience | |
| CD115 | AFS98 | APC | eBioscience | |
| c-kit | 2B8 | APC-Cy7 | BioLegend | |
| Sca1 | D7 | BUV395 | BD Biosciences | |
| CD127 | A7-R34 | PE-Dazzle594 | BioLegend | |
| Flt3 | A2F10 | BV421 | BioLegend | |
| Ly6D | 49-H4 | PE | BioLegend | |
| CD317 | ebio927 | Alexa Fluor 488 | eBioscience | |

#### Figure 3B-C : Spleen cDC phenotypic analysis

| Specificity | Clone | Fluorochrome or biotin | Source |
|---|---|---|---|
| CD16/32 | 2.4G2 | - | BD Biosciences |
| CD11c | HL3 | PE-Cy7 | BD Biosciences |
| CD8a | 53-6.7 | PerCP-Cy5.5 | BD Biosciences |
| CD11b | M1/70 | APC-Cy7 | BD Biosciences |
| I-A$^b$ (MHC II) | AF6-120.1 | eFluor 450 | eBioscience |
| H-2K$^b$ (MHC I) | AF6-88.5 | Alexa Fluor 647 | BD Biosciences |
| CD80 (B7-1) | 16-10A1 | PE | BD Biosciences |
| CD86 (B7-2) | GL1 | FITC | BD Biosciences |

#### Figure 3D-H : Spleen cDC FACS-sorting for epigenetic analysis and in vitro culture

| Specificity | Clone | Fluorochrome or biotin | Source |
|---|---|---|---|
| CD16/32 | 2.4G2 | - | BD Biosciences |
| CD11c | HL3 | PE | BD Biosciences |
| I-A$^b$ (MHC II) | AF6-120.1 | APC | eBioscience |

##### Figure 3I : Spleen B cell, pDC, cDC1 and cDC2 FACS-sorting for RT-qPCR analysis

| Specificity | Clone | Fluorochrome or biotin | Source |
|---|---|---|---|
| CD16/32 | 2.4G2 | - | BD Biosciences |
| CD11c | HL3 | PE | BD |
| CD19 | 1D3 | APC | BD |
| CD45R (B220) | RA3-6B2 | PerCP-Cy5.5 | BD |
| I-A$^b$ (MHC II) | AF6-120.1 | FITC | BD |
| CD11b | M1/70 | APC-Cy7 | BD |
| CD8a | 53-6.7 | PE-Cy7 | BD |

##### Figure 5 : T cell FACS-sorting and analysis

**CD4$^+$ T cells FACS-sorting from OT-II. *Ly5.1.Foxp3EGFP* mice**

| Specificity | Clone | Fluorochrome or biotin | Source |
|---|---|---|---|
| CD16/32 | 2.4G2 | - | BD Biosciences |
| CD4 | RM4-5 | PE-Cy7 | BD Biosciences |

**CD4$^+$ and CD8$^+$ T cell proliferation analysis (dilution of eFluor 670 CPD)**

| Specificity | Clone | Fluorochrome or biotin | Source |
|---|---|---|---|
| CD16/32 | 2.4G2 | - | BD Biosciences |
| CD45.1 | A20 | APC-Cy7 | BD Biosciences |
| CD45.2 | 104 | PE | BD Biosciences |
| CD4 | RM4-5 | V450 | BD Biosciences |
| CD8a | 53-6.7 | PE-Cy7 | BD Biosciences |

##### Figure 6B : Endocytosis/degradation assays

| Specificity | Clone | Fluorochrome or biotin | Source |
|---|---|---|---|
| CD16/32 | 2.4G2 | - | BD Biosciences |
| CD11c | HL3 | PE-Cy7 | BD Biosciences |
| CD8a | 53-6.7 | PerCP-Cy5.5 | BD Biosciences |
| CD11b | M1/70 | APC-Cy7 | BD Biosciences |
| I-A$^b$ (MHC II) | AF6-120.1 | eFluor 450 | eBioscience |

##### Figure 6C : Surface or total (intra) expression of MHC II-associated molecules

| Specificity | Clone | Fluorochrome or biotin | Source |
|---|---|---|---|
| CD16/32 | 2.4G2 | - | BD Biosciences |
| CD45R (B220) | RA3-6B2 | PerCP-Cy5.5 | BD Biosciences |
| CD11c | N418 | BV421 | BD Biosciences |
| CD11bA:BA:CA:DA:EA:D | M1/70 | APC-Cy7 | BD Biosciences |
| CD8a | 53-6.7 | PE-Cy7 | BD Biosciences |
| I-A$^b$ (MHC II) | AF6-120.1 | FITC | BD Biosciences |
| H2-O(b) | Mags.Ob1 | Alexa Fluor 647 | Lisa Denzin |
| H2-M(αβ2) | 2E5A | Purified | BD Biosciences |
| Anti rat IgG1 (for H2-M) | RG11/39.4 | FITC | BD Biosciences |
| CD74 | In-1 | FITC | BD Biosciences |
| I-Ab-CLIP | 15G4 | FITC | Santa Cruz |

#### Oligonucleotides used in this study

##### Primers used for qPCR analysis of *C.Rodentium* (genomic DNA template)

| Gene name | Forward primer sequence (5' to 3') | Reverse primer sequence (5' to 3') | Amplicon size (bp) |
|---|---|---|---|
| *EspB* (*C.Rodentium*)[a] | ATGCCGCAGATGAGACAGTTG | CGTCAGCAGCCTTTTCAGCTA | 141 |
| *16S* rRNA total bacteria | GGTGAATACGTTCCCGG | TACGGCTACCTTGTTACGACTT | 144 |

[a] Sagaidak, S., Taibi, A., Wen, B. & Comelli, E. M. Development of a real-time PCR assay for quantification of Citrobacter rodentium. J. Microbiol. Methods 126, 76–77 (2016)

##### Primers used for RT-qPCR analysis (cDNA template)

| Gene name | Forward primer sequence (5' to 3') | Reverse primer sequence (5' to 3') | Amplicon size (bp) |
|---|---|---|---|
| *Tmem176a* | CAAACTTCTGCTGGCCGGAT | GTGAAGGAAGGCAACAGCTC | 238 |
| *Tmem176b* | AAGAAGTTTCTCTCCTGGCCT | CAGTTTTCCCTGCCTCTTCTCA | 222 |
| *Rorc2* | GGAGGACAGGGAGCCAAGTT | AGTAGGCCACATTACACTGCT | 160 |
| *Il23r* | GCAACATGACATGCACCTGG | TGTTGTGAGTTCTCCATGCCT | 208 |
| *Il22* | CCTACATGCAGGAGGTGGTG | AAACAGCAGGTCCAGTTCCC | 176 |
| *Il17a* | AGTCCAGGGAGAGCTTCATCT | TCTTCATTGCGGTGGAGAGTC | 248 |
| *Il17f* | GAAGTGCACCCGTGAAACAG | AACTGGAGCGGTTCTGGAAT | 230 |
| *Reg3g* | CAGATATGGCCTGCCAAAAGA | TGTTGGGTTCATAGCCCAGTG | 159 |
| *Defb2* | CTGGAGTCTGAGTGCCCTTTC | CTCCATTGGTGTGGCAGTGG | 153 |
| *Lcn2* | ATGTCACCTCCATCCTGGTC | GCGAACTGGTTGTAGTCCGT | 169 |
| *S100a7a* | CAGACACACCAGTGGAGGAC | ATGTAGTATGGCTGCCTGCG | 169 |
| *Cxcl3* | CCCAGACAGAAGTCATAGCCAC | CCCAGACAGAAGTCATAGCCAC | 193 |
| *Cxcl5* | GCTGGCATTTCTGTTGCTGT | AGTTTAGCTATGACTTCCACCGT | 192 |
| *Gapdh* | GGTGAAGGTCGGTGTGAACGG | TCGCTCCTGGAAGATGGTGAT | 232 |
| Cre | CGACCAGGTTCGTTCACTCA | CAGCGTTTTCGTTCTGCCAA | 184 |

##### Primers used for *Tmem176a/b* germline DKO mouse genotyping (genomic DNA template)

| Gene | Primer name | Sequence (5' to 3') |
|---|---|---|
| *Tmem176a* | AF | CAGCATCAGGCTTCCCTGAC |
| *Tmem176a* | AR | GCCACACTCACCACGATCCC |
| *Tmem176b* | BF | TGAAACTCTTCTCCTGTCACCCA |
| *Tmem176b* | BR | TAATCCTTTGGGGAGGGGGCC |

| Primer combination[b] | Expected amplicon sizes (bp)<br>*Tmem176a/b* WT mouse (+/+) | DKO mouse (–/–) | Heterozygous mouse (+/–) |
|---|---|---|---|
| AF + AR | 667 | - | 667 |
| BF + BR | 620 | - | 620 |
| BF + AR | 4769[c] | 572 | 4769[c] + 572 |
| AF + AR + BF | 4769[c] + 667 | 572 | 4769[c] + 667 + 572 |
| BF + BR + AR | 4769[c] + 620 | 572 | 4769[c] + 620 + 572 |

+ : WT allele, – : DKO allele (large deletion)

[b] We routinely use the 3-primer combination AF+AR+BF

[c] No amplification of this large amplicon using short elongation times (e.g. 20 s) during PCR

##### Primers used for *Tmem176a/b* conditional (« floxed ») DKO mouse genotyping (genomic DNA template)

| Gene | Primer name | Sequence (5' to 3') |
|---|---|---|
| *Tmem176a* | aEf | ATGGCGGTCCCTTTGCTTCC |
| *Tmem176a* | aEr | TGCACTCCGCTTTCCCTGTCTT |
| *Tmem176b* | bEf | CTGTTCCTGGCACTCCTTTACCTTGG |
| *Tmem176b* | bEr | GCTGCAGATGCAAGACAACACTAACCA |

| Primer combination[d] | Expected amplicon sizes (bp)<br>*Tmem176a/b* WT mouse (wt/wt) | Homozygous floxed mouse (flox/flox)[d] | Heterozygous floxed mouse (wt/flox) |
|---|---|---|---|
| aEf + aEr | 280 | 402 | 280 + 402 |
| bEf + bEr | 238 | 386 | 238 + 386 |

[d] These combinations distinguish between WT and intact floxed alleles. To detect null allele resulting from Cre-mediating recombination, use bEf + aEr : null = 427 bp, flox = 5640 bp, WT = 5370 bp

##### Primers used for Cre-transgenic mouse genotyping (genomic DNA template)

| Gene | Primer name | Sequence (5' to 3') |
|---|---|---|
| Cre | Cre-F | CGACCAGGTTCGTTCACTCA |
| Cre | Cre-R | CAGCGTTTTCGTTCTGCCAA |
| Internal control *18S* | 18S-F | AGTTCCGACCATAAACGATGC |
| Internal control *18S* | 18S-R | CCCTTCCGTCAATTCCTTTAA |

| Primer combination[d] | Expected amplicon sizes (bp)<br>Cre-negative mouse | Cre-positive mouse[e] |
|---|---|---|
| Cre-F + Cre-R | - | 184 |
| 18S-F + 18S-R | 143 | 143 |

[e] This assay will not distinguish heterozygous from homozygous Cre-transgenic mice. We maintained our Cre-expressing mice in heterozygosity

#### Plasmids used in this study

##### RUSH constructs

| Protein of interest (Uniprot#) | Full name of the plasmid | Fluorescent protein / Hook | Source |
|---|---|---|---|
| TMEM176B (Q3YBM2) | Str-Ii_**TMEM176B**-SBP-**EGFP** | EGFP / Ii | This study |
| TMEM176B (Q3YBM2) | Str-Ii_**TMEM176B**-SBP-**mCherry** | mCherry / Ii | This study |
| TMEM176B (Q3YBM2) | Str-G84_**TMEM176B**-SBP-**mCherry** | mCherry / Golgin-84 | This study |
| TMEM176A (Q96HP8) | Str-Ii_**TMEM176A**-SBP-**EGFP** | EGFP / Ii | This study |
| Golgin-84 (Q8TBA6 ) | Str-Ii_SBP-**EGFP**-**Golgin-84** | EGFP / Ii | Franck Perez - Boncompain *et al.*[1] |
| TNFa (P01375) | Str-KDEL_**TNFa**-SBP-**EGFP** | EGFP / KDEL | Franck Perez - Boncompain *et al.*[1] |
| CD44 (P16070) | Str-Ii_**CD44**-SBP-**EGFP** | EGFP / Ii | Franck Perez - Boncompain *et al.*[1] |
| LAMP1 (P11279) | Str-Ii_**LAMP1**-SBP-**mCherry** | mCherry / Ii | This study |
| TfR (P02786) | Str-KDEL_**TfR**-SBP-**EGFP** | EGFP / KDEL | Juan S. Bonifacino - Chen *et al.*[2] |
| CD-MPR (P20645) | Str-KDEL_SP-SBP-**EGFP**-**CD-MPR** | EGFP / KDEL | Juan S. Bonifacino - Chen *et al.*[2] |

Str : Core streptavidin

Ii : Isoform of human invariant chain retained in the ER, used as a hook

G84 : Golgin-84, resident of the Golgi apparatus, used as a hook

KDEL : ER retention signal (Lysine [K], Aspartic acid[D], Glutamic acid [E], Leucine [L]), used as a hook

SBP : Streptavidin-binding peptide

##### Other constructs

| Protein of interest | Full name of the plasmid | Fluorescent protein | Source |
|---|---|---|---|
| GGA1 | GFP-**GGA1** | EGFP | Juan S. Bonifacino - Puertollano *et al.*[3] |
| RAB4B | GFP-**RAB4B** | EGFP | Chen *et al.*[4] (provided by Juan S. Bonifacino) |
| RAB5(Q79L) | mCherry-**RAB5CA(Q79L)** | mCherry | Addgene#35138, gift from Sergio Grinstein - Bohdanowicz *et al.*[5] |
| RAB7A | GFP-**RAB7A** | EGFP | Juan S. Bonifacino - Rojas *et al.*[6] |
| HLA-DMα | DMb-**DMa**-YFP | YFP | Jacques Neefjes - Zwart *et al.*[7] |
| HLA-DRβ | DRa-**DRb**-YFP | YFP | This study (using the DRa-DRb-CFP construct from Zwart *et al.*[7]) |
